# Multifunction Fluorescence Open-Source *In Vivo/In Vitro* Imaging System (openIVIS)

**DOI:** 10.1101/2023.10.06.561111

**Authors:** John M. Branning, Kealy A. Faughnan, Austin A. Tomson, Grant J. Bell, Sydney M. Isbell, Allen DeGroot, Lydia Jameson, Kramer Kilroy, Michael Smith, Robert Smith, Landon Mottel, Elizabeth G. Branning, Zoe Worrall, Frances Anderson, Ashrit Panditaradyula, William Yang, Joseph Abdelmalek, Joshua Brake, Kevin J. Cash

## Abstract

The widespread availability and diversity of open-source microcontrollers paired with off-the-shelf electronics and 3D printed technology has led to the creation of a wide range of low-cost scientific instruments, including microscopes, spectrometers, sensors, data loggers, and other tools that can be used for research, education, and experimentation. These devices can be used to explore a wide range of scientific topics, from biology and chemistry to physics and engineering. In this study we designed and built a multifunction fluorescent open-source in-vivo/in-vitro imaging system (openIVIS) fluorescent imaging system that integrates a Raspberry Pi with commercial cameras and LEDs with 3D printed structures combined with an acrylic housing. Our openIVIS provides three excitation wavelengths of 460 nm, 520 nm, and 630 nm integrated with Python control software to enable fluorescent measurements across the full visible light spectrum. To demonstrate the various potential applications of our system, we tested its performance against a diverse set of experiments including laboratory type assays (measuring fluorescent dyes, using optical nanosensors, and DNA gel electrophoresis) to potentially fieldable applications (plant and mineral imaging). We also tested the potential use for a high school biology environment by imaging small animals and tracking their development over the course of a couple of weeks. Our system demonstrated its ability to measure a wide dynamic range fluorescent response from millimolar to picomolar concentrations in the same sample while measuring responses across visible wavelengths. These results demonstrate the power and flexibility of open-source hardware and software and how it can be integrated with customizable manufacturing to create low-cost scientific instruments with a wide range of applications. Our study provides a promising model for the development of low-cost instruments that can be used in both research and education.

## Introduction

Over the decades biological imaging has become an integral part of research in biological, medical, and chemical sciences. While there are a wide range of modalities used in research such as magnetic resonance imaging (MRI), positron emission tomography (PET), and computed tomography (CT), optical fluorescence imaging is a key player in early-stage research across academia and industry. Imaging systems coupled with fluorescent dyes and high-resolution cameras allow researchers to investigate cellular processes and dynamics non-invasively in two and three dimensions (1). This has enabled scientists to quantify metabolite concentrations, track molecular interactions, and observe tissues from subcellular structures and cells to the entire organism (2,3). However, the high cost associated with commercial fluorescent imaging systems has precluded the proliferation of imaging to smaller laboratories and research organizations with limited research budgets. The emergence of free open-source software (FOSS) and free open-source hardware (FOSH) coupled with three-dimensional (3D) printing technology and other rapid prototyping approaches has encouraged the development of low-cost laboratory and scientific equipment (4–7). The proliferation of FOSS and FOSH allows and encourages developers and students from non-traditional environments like high school chemistry and biology classes to engage in scientific research. In addition, the capabilities of commercially available cameras have increased drastically while the cost per pixel has plummeted allowing open-source designs to approach commercial scientific equipment capabilities with cost savings approaching 94% of the commercial equipment while at the same time increasing reproducibility in lab equipment (6,8). These developments have spurred research groups to develop their own low-cost laboratory equipment for various applications. Several research groups have developed open-source fluorescence imaging systems that were able to monitor the growth of bacterial colonies on agar plates (9,10). Other researchers designed and built Raspberry Pi based microscope systems capable of imaging cellular structures (11–13). Researchers also fabricated open-source plate readers to supplement commercial plate readers for measuring absorbance and fluorescence emissions using standard laboratory well plates (14–16). There are many other examples of open-source equipment that researchers have developed that are outside the scope of this paper that interested readers can use for further inspiration for their own designs (6).

*In Vivo* Imaging Systems (IVIS) refer to a group of optical imaging instruments used to visualize and study biological processes inside a living organism in real-time. These systems are designed to non-invasively monitor various physiological and pathological changes that occur within living systems via fluorescent and bioluminescent responses. Typical IVIS applications range from studying the progression of infectious diseases, drug pharmacokinetics, oncology studies, to imaging transgenic small animals. Commercial optical IVIS systems are available from several companies including PerkinElmer (17), Spectral Instruments Imaging (18), LI-COR Biosciences (19), and Azure Biosciences (20). These systems typically consist of a cooled charge couple device (CCD) camera, an optical filter wheel, multiple excitation sources, optical lenses, a sample stage, and control electronics. Commercial systems offer a large range of excitation and filter wavelengths to enable researchers to use a wide range of fluorescent and bioluminescent dyes in their experiments. These systems are coupled to extremely sensitive imagers to enable the long exposure times necessary for bioluminescent imaging. However, the high performance of the commercial systems comes with a high price point with some systems costing several hundred thousand dollars and requiring proprietary control and analysis software. Several research groups have developed low-cost open source IVIS systems for use in the laboratory and classroom (9,10,21). While the open-source systems offer reduced performance in terms of excitation and filter wavelength options, imaging sensitivity, and sample size when compared to the commercial systems, they enable more researchers the ability to leverage *in vivo* imaging for their research democratizing access to this imaging modality.

In our work, we combine low-cost cameras, open-source microelectronics, glass filters, acrylic housing, 3D printed housings, and off-the-shelf light-emitting diodes (LEDs) and control electronics to create an integrated multifunction fluorescent open-source *in-vivo/in-vitro* imaging system (openIVIS) using three excitation wavelengths and emission wavelengths only limited by filter availability. The system was designed to track metabolite and ion levels via fluorescent dyes in standard laboratory well plates as well as *in vivo* in small animals and other biological systems with sizes ranging from 250-500 μm for a single algae or bacteria colony to 150 mm for small animals. Our system (Fig 1) was validated through a series of experiments including animals, plants, and minerals (everything from the Linnaean taxonomy (22)) to show the wide range of applications for use in science, technology, engineering, and mathematics (STEM) education, from secondary to collegiate levels, as well as scientific and engineering research.

**Fig 1.**
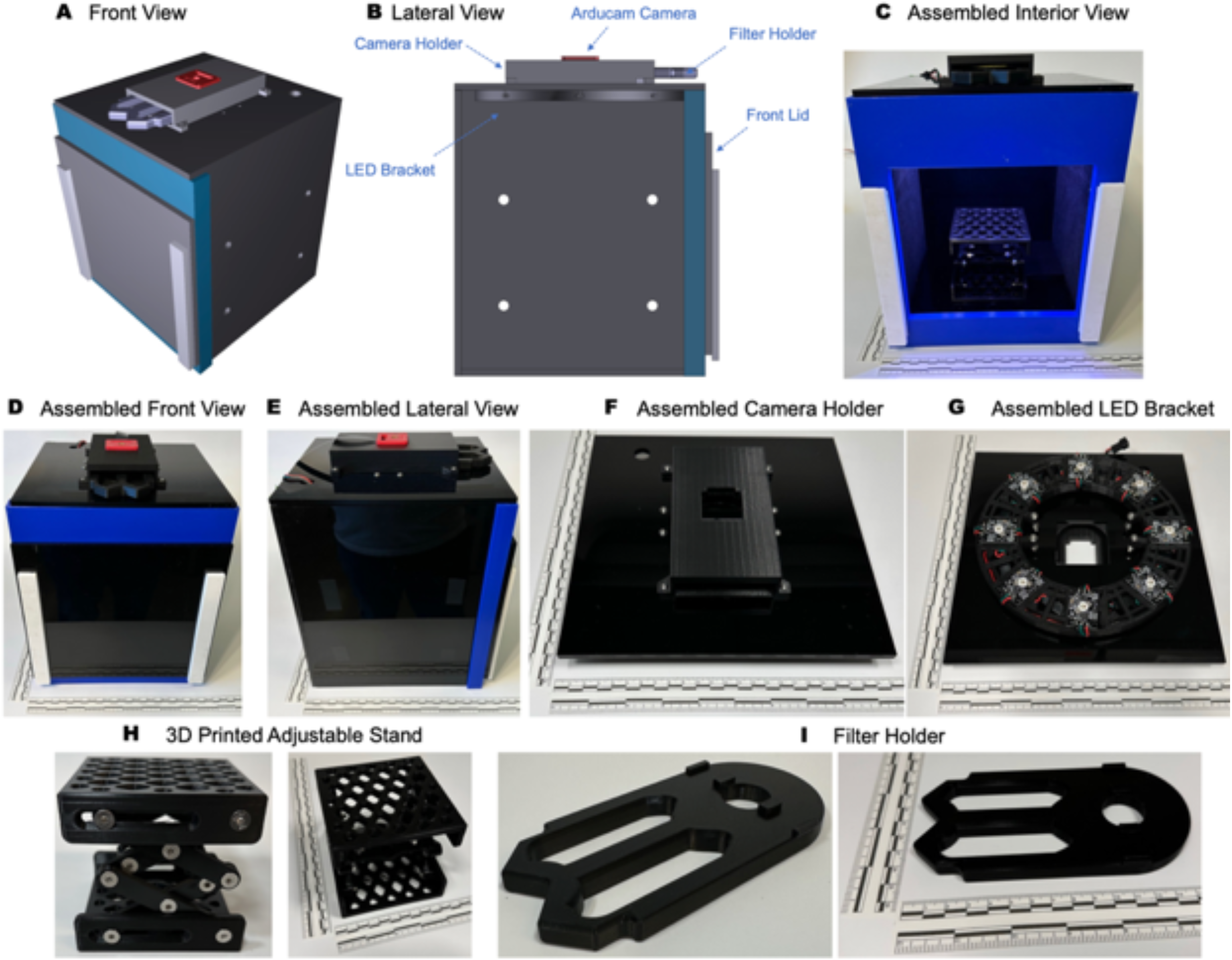
openIVIS System Architecture. (A) 3D rendering of imaging system, (B) 3D rendering lateral view of a longitudinal cross-section imaging box (C) Front view of imaging box with front cover removed showing 3D printed adjustable stand for holding specimens (D) Front view of assembled imaging system, (E) Lateral view of assembled imaging system, (F) Assembled camera holder, (G) Assembled LED bracket with 3 color LED, (H) Adjustable stand for holding samples, (I) Removable filter holder Higher resolution images for all panels are available in SI.

## Materials and Methods

### Imaging Box

The components that hold the camera, optical filters, excitation LEDs, adjustable stand and sample holder were fabricated via 3D printing using an Original Prusa i3 MK3S+ printer using 1.75 mm polyactic acid (PLA) 3D printer filament (Amazon, Part Number: B00J0ECR5I). A color calibration (B00AWT2QCE) and lens test chart (B00F1YEEHA) were purchased from Amazon. Unmounted twenty-five milimeter long pass optical glass filters (FGL435, FGL455, FGL495, FGL515, FGL530, FGL550, FGL570, FGL590, FGL610, FGL630, FGL645, FGL665, FLG695) were purchased from ThorLabs. A bill of materials (Table 1 in S1) for the control electronics is available in the supporting information.

### Nanosensor Fabrication

Poly(vinyl chloride) (high molecular weight, (PVC)), tetrahydrofuran (THF), dichloromethane (DCM), sodium tetrakis[3,5-bis(trifluoromethyl)phenyl]borate (NaBARF; Selectophore), bis(2-ethylhexyl) sebacate (DOS/BEHS, Selectophore), 9-dimethylamino-5-[4-(15-butyl-1,13-dioxo-2,14-dioxanonadecyl)phenylimino]benzo[a]phenoxazine (chromoionophore VII; CHVII), 6,6-dibenzyl-1,4,8,11-tetraoxacyclotetradecane (lithium ionophore VI; LiI VI), trioctylphosphine oxide (TOPO), tetrahydrofuran (THF), 4-(2-hydroxyethyl)piperazine-1-ethanesulfonic acid (HEPES), dichloromethane (DCM), 9-dimethylamino-5-[4-(16-butyl-2,14-dioxo-3,15-dioxaeicosyl)-phenylimino]benzo[a]phenoxazine (chromoionophore II; CHII), N,N-dicyclohexyl-N0,N0-diisobutyl-cis-cyclohexane-1,2-dicarboxamide (lithium ionophore III; LiI III), 2-nitrophenyl octyl ether (NPOE), and lithium chloride were purchased from Sigma-Aldrich (St. Louis, MO, USA). 1,2-Dipalmitoyl-sn-glycero-3-phosphoethanolamine-N-[methoxy(polyethylene glycol)-750] ammonium salt in chloroform (PEG-lipid) was purchased from Avanti Polar Lipids (Alabaster, AL). 2-[4-(2-Hydroxyethyl)piperazin-1-yl]ethanesulfonic acid (HEPES; Molecular Biology grade), 2-amino-2-hydroxymethylpropane-1,3-diol (Tris; 2 M), hydrochloric acid concentrate (HCl; 10 N, ACS certified), sodium hydroxide concentrate (NaOH; 10 N, ACS certified), 2-Amino-2-hydroxymethylpropane-1,3-diol, 2M solution (TRIS, 2M), and fluorescein-5-isothiocyanate (FITC) was purchased from Fisher Scientific (Waltham, MA, USA). 1,10-Dioctadecyl-3,3,30,30-tetramethylindotricarbocyanine iodide (DiR) was purchased from Life Technologies (Grand Island,NY,USA). Phosphate-buffered saline (PBS with Ca2þ and Mg2þ, pH = 7.4) was purchased from Boston Bioproducts (Ashland, MA, USA). Lithium and sodium nanosensor were fabricated in HEPES-TRIS buffer using established protocols (23,24) and concentrated to 10× using a centrifuge at 14,000 RPM for 14 minutes.

### Other Chemicals

A 1 kb DNA Ladder ready-to-use (ThermoFisher, 100 ng/µL) was used as DNA ladder marker. Molecular biology agarose (Bio-Rad) was used for DNA gel electrophoresis. Invitrogen SYBR Safe DNA Gel Stain (ThermoFisher) supplied as a 10,000× concentrate in DMSO was used as loading dye. TAE buffer (BIO-RAD) supplied as a 50× concentrate was used as running buffer. DNA agarose gel electrophoresis was performed for 50 min at 120 V with 1.0% agarose gel in a 1× TAE buffer (25). Tobacco hornworm, *Manduca sexta*, larvae were purchased from Carolina. A fluorescent mineral kit was purchased from Frey Scientific. Organic green leaf lettuce was purchased from a local Golden, CO grocery store.

### Instrumentation

Detailed information on system operation is included in supporting information, S5 and S6, and is also available online (https://github.com/CashLabMines/openIVIS.git). For Fig 2, Fig 3, and Fig 8 the camera was operated using the default exposures of libcamera-still. For Fig 5, Fig 6, Fig 7, Fig 9, and Fig 10 the camera was operated using libcamera-still while looping through shutter times from 1-10000 to adjust the camera exposure time in milliseconds. The LEDs were controlled using the NeoPixel library in python using the RGB color code values for the blue LED (460 nm), green LED (520 nm) and red LED (630 nm) excitations. For external excitation such as UV the pixel color code was set to zero. To change the LED power level the pixel brightness was changed to a value between 0 – 1.0 based upon the desired output power.

**Fig 2.**
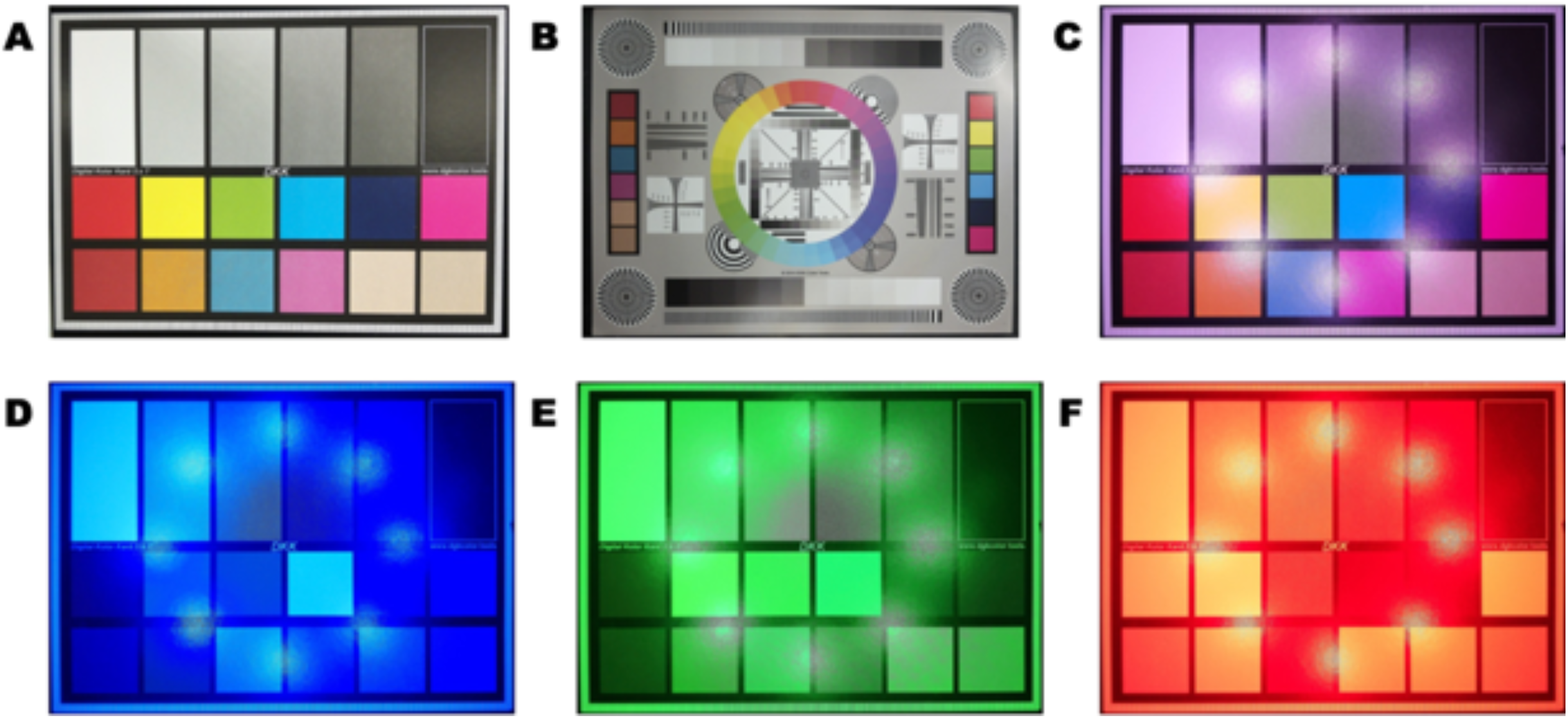
Camera Calibration. (A) Image of color checker card used for color verification (B) Image of photo calibration chart show lines and patterns for distortion analysis, white and black levels (C) Image of color checker under unbalanced white LED excitation (D) Image of color checker under blue LED (460 nm) excitation (E) Image of color checker under green LED (520 nm) excitation (F) Image of color checker under blue LED (630 nm) excitation

**Fig 3.**
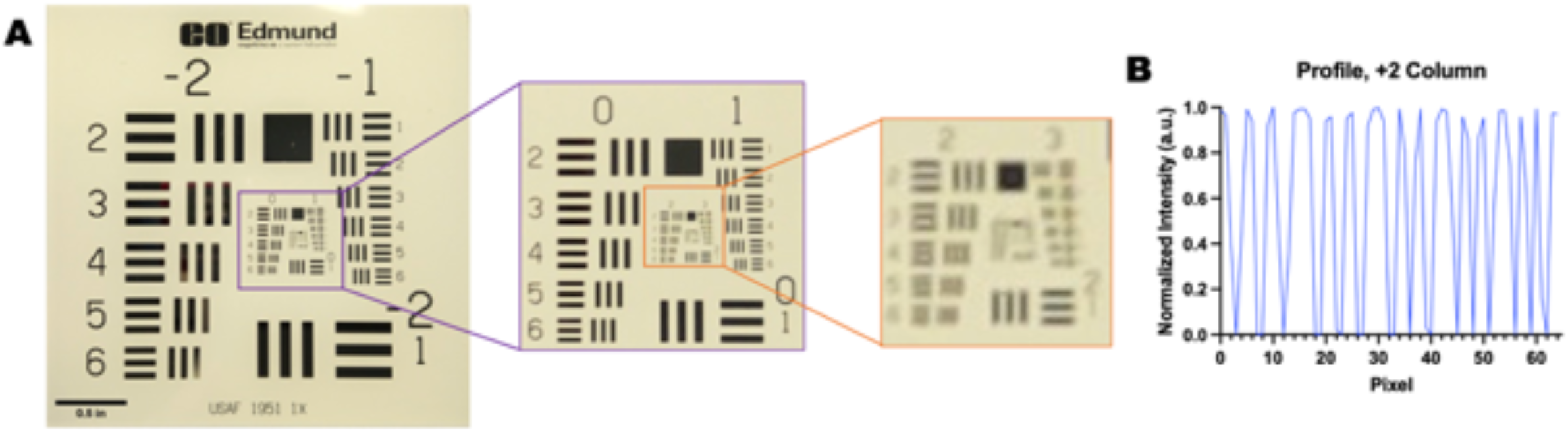
openIVIS Resolution Portion of a single image under full spectrum illumination showing a 1951 U.S. Air Force resolution target. The image indicates system has a resolution greater than 6-line pairs/mm which is equivalent to 79 μm.

**Fig 4.**
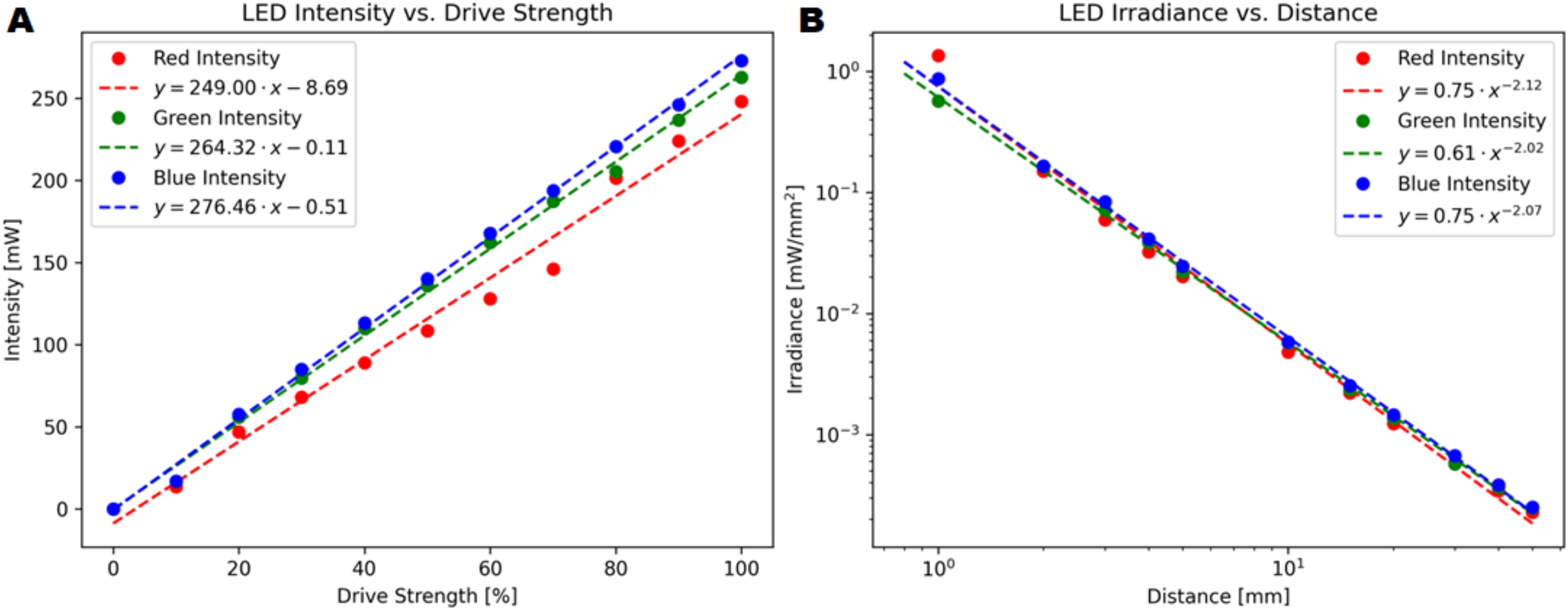
LED excitation power characterization. (A) Measured power linearity versus drive strength (B) Measured power at varying distances

**Fig 5.**
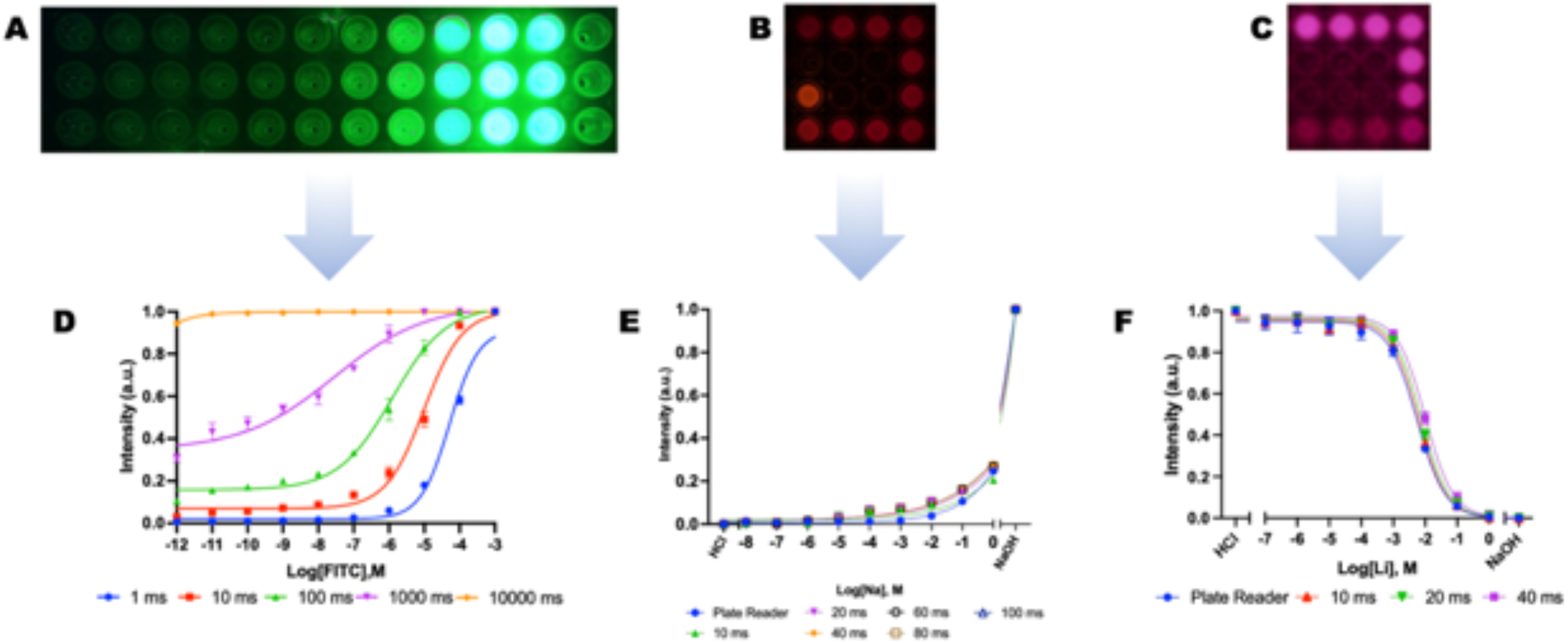
Imaging of fluorescent nanosensors. (A) Subimage of a FITC serial dilution in 96 well PCR plate with a 100 ms exposure (B) Sub-image of sodium nanosensors in the center of a 96-well PCR plate with a 80 ms exposure (C) Sub-image of lithium nanosensors in the center of a 96-well PCR plate with a 40 ms exposure (D) Normalized FITC Sensitivity curves versus exposure time, n = 3 (E) Normalized sodium nanosensor characterization with green (520 nm) excitation with long-pass 590 nm optical filter, n = 3 (F) Normalized Lithium nanosensor characterization with red (630 nm) excitation with long-pass 665 nm optical filter, n = 3

**Fig 6.**
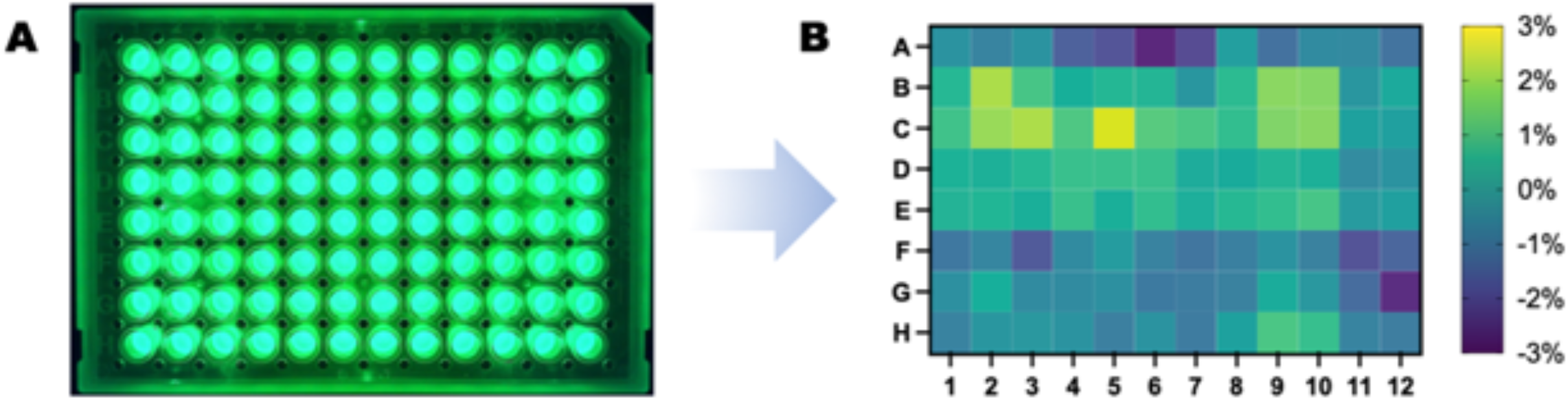
Imaging System fluorescence flatness from a 96 well plate with 10 µM of FITC in each well with a blue LED (465 nm) excitation and a bandpass 542 nm ± 25 nm optical glass filter. (A) Image of 96 well plate with 10 µM of FITC (B) Percent deviation of each well from plate average, n = 3

**Fig 7.**
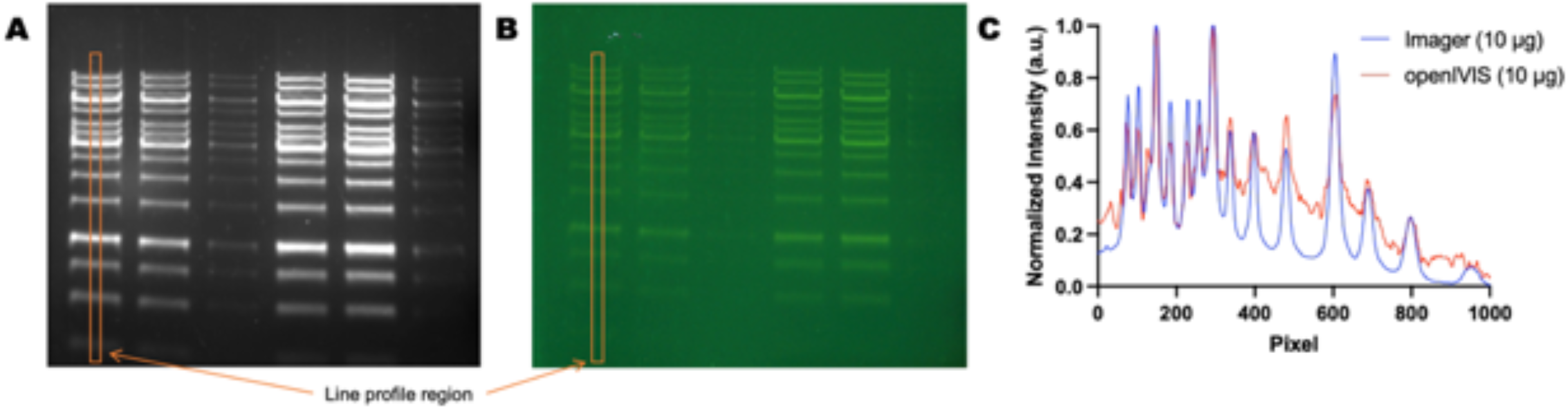
Detection of DNA ladder makers stained with SYBR Safe. (A) Detection using Bio-Rad ChemiDoc MP Imaging System (B) Detection by blue LED light (465 nm) excitation with long pass 530 nm optical glass filter (C) Line profile through left 10 µg lane in each image.

**Fig 8.**
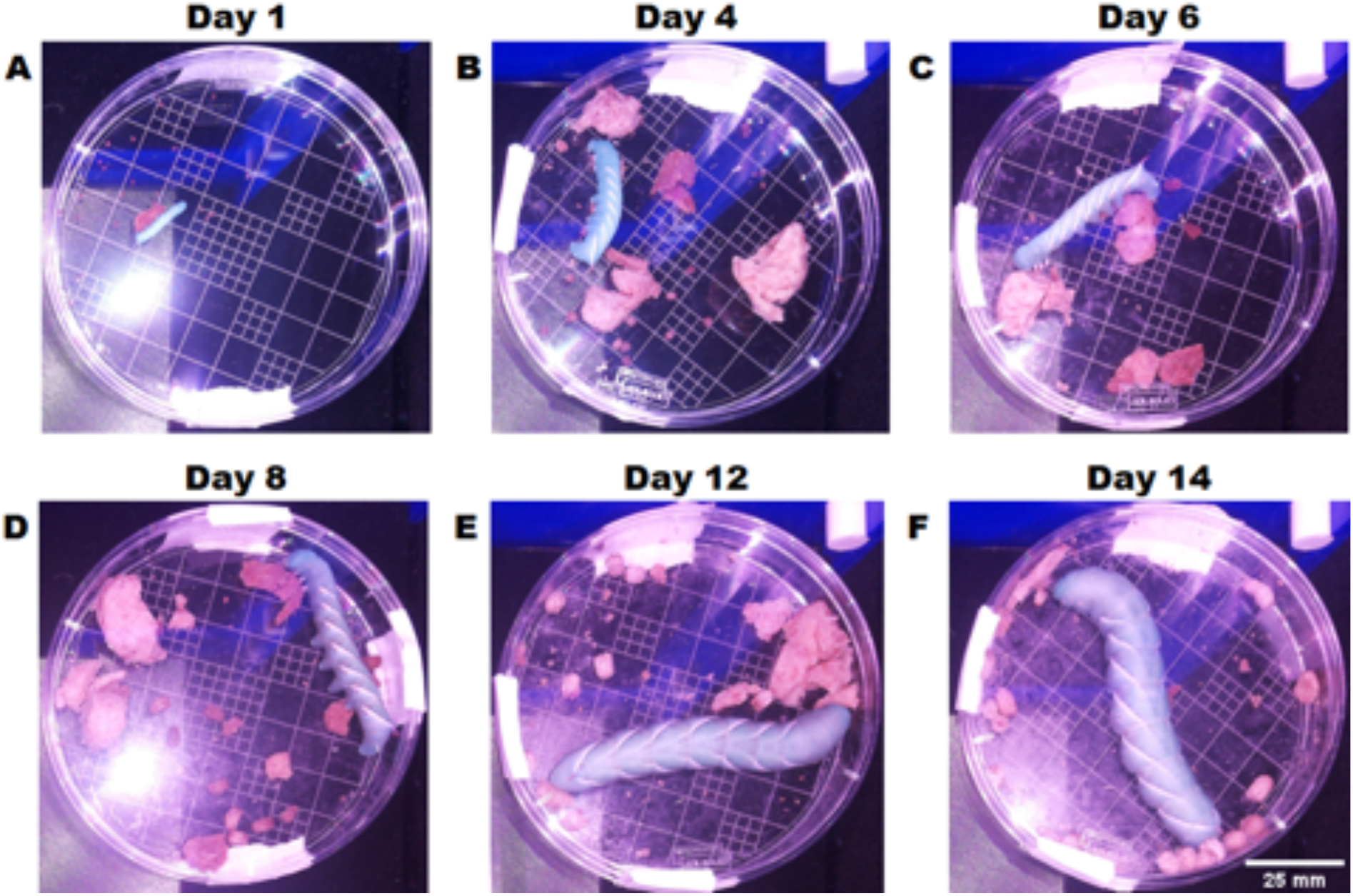
Monitoring growth of hornworms over time.

**Fig 9.**
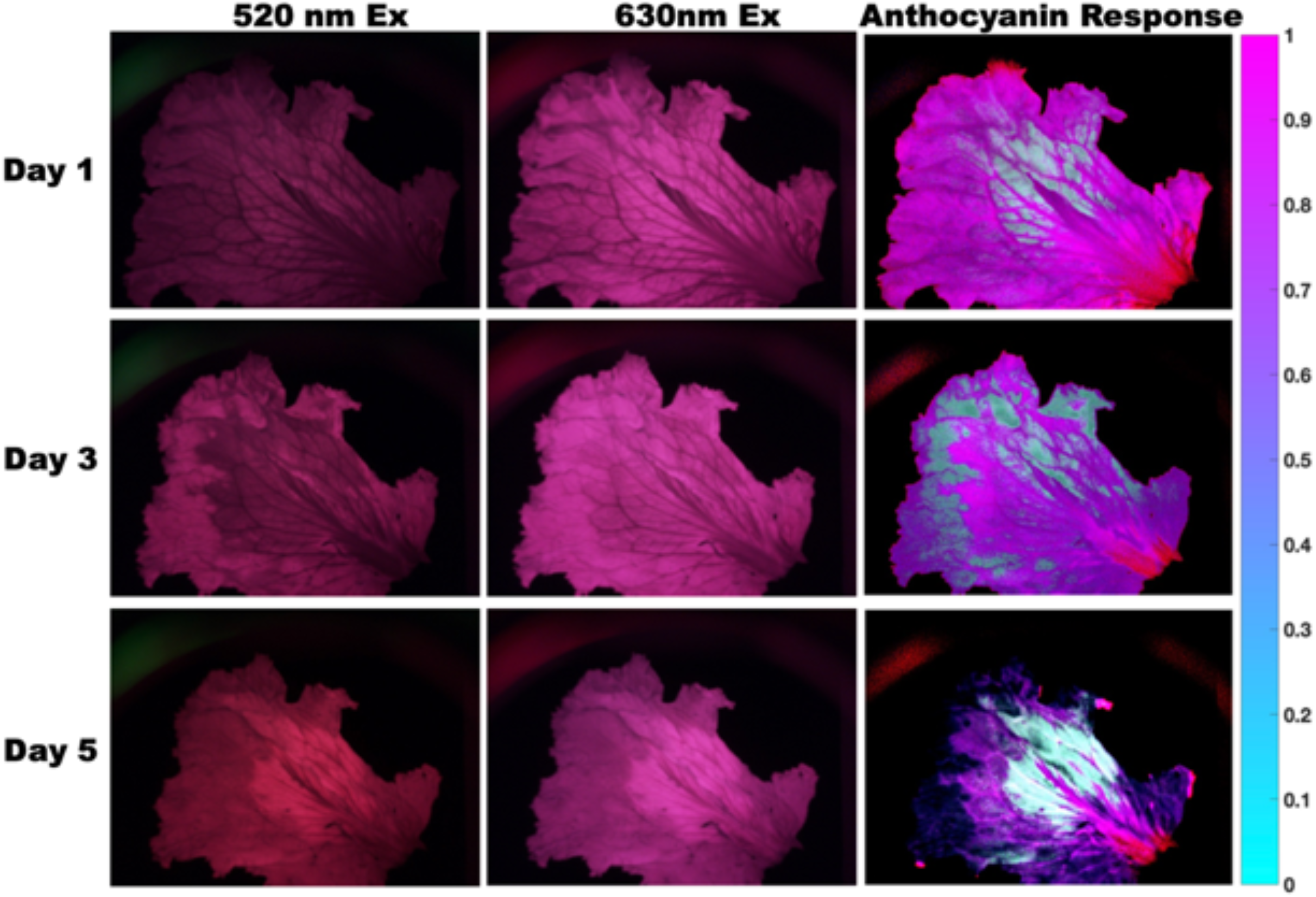
Images of lettuce leaf decaying over time. For each row the left column shows 520 nm excitation with 665 nm long pass filter, center column shows 630 nm excitation with 665 nm long pass filter, right column shows measured anthocyanin response.

**Fig 10.**
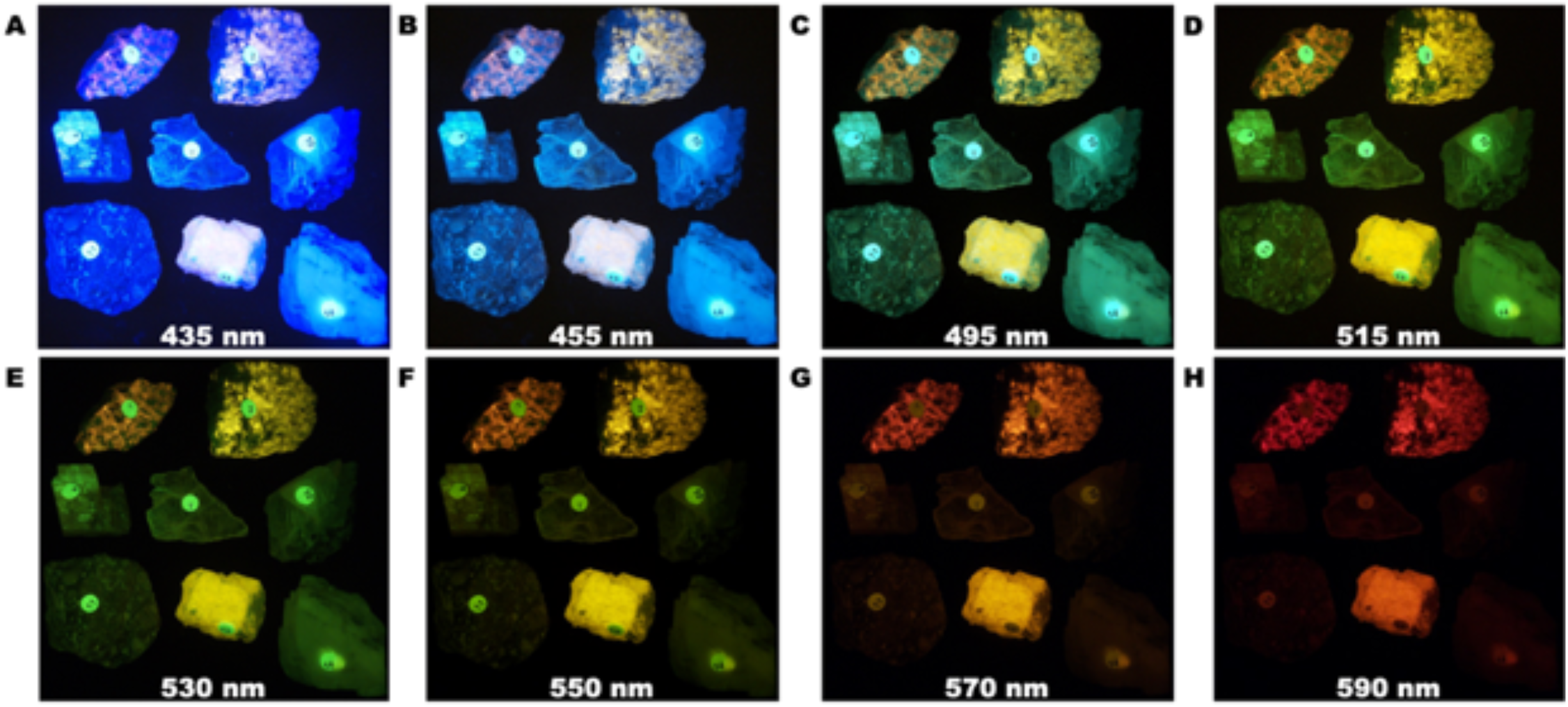
Fluorescence response of various minerals excited with long wave (395 nm) UV light imaged with varying long pass optical glass filters (A-H)

### Results and Discussion System Design Decisions Imaging Box

We designed an open-source fluorescent imaging system comprised of low-cost cameras, open-source microelectronics, glass filters, acrylic housing, 3D printed housings, and off-the-shelf light-emitting diodes (LEDs) and control electronics. Fig 1 shows a system diagram of the assembled imaging system with further diagrams, supporting computer aided design (CAD), and 3D printer standard triangle language (STL) files available in the supporting information (S1). The imaging housing was fabricated from black acrylic sheets laser cut to the specified dimensions. The main body of the box consists of 6 acrylic sides, with one of them containing a large hole in the front to allow for the placing of specimens into the box. The front of the box is covered with another acrylic piece that slides over the 3D print at the front of the box to hold it in place. The inside walls of the box were covered with adhesive backed black felt fabric sheets to reduce the internal reflection from the acrylic. The sides of the box have holes cut for optional shelfs. The top acrylic piece of the box has holes for screwing the filter and camera holder 3D print to the top of the box and a place for wiring from the LEDs to exit the box.

The LED holder, Fig 1G, was designed to equally space the eight LEDs around the camera equidistance from the center point of the box to the edge to provide uniform LED illumination across the imaging samples. An 3D printed adjustable stand, Fig 1H, was designed and printed to allow for adjusting the height of the imaging samples withing the imaging box. The sample holder and adjustable stand had a preset slot that centered the samples in the camera field of view (FOV) and LED illumination area. In our imaging system we used glass optical filters to remove the unwanted fluorescence emission wavelengths (dichroic filters also work but were not chosen for this application due to costs). We designed a simple 3D printed filter holder, Fig 1I, that allowed us to quickly swap out optical filters as needed throughout out experiments without disturbing any other parts of the system. The benefit of using 3D printed technology is that we could easily print out a different holder to accommodate bigger or smaller filter sizes, and the filter holder was designed to enable filters as large as 27 mm without needing to redesign any other parts of the system, and with minor changes to the filter holder the system could accommodate larger filters as well. The flexibility of the filter holder allows the end user to quickly swap out filters enabling the user to use a wide range of filter wavelengths based on their needs. The flexibility of 3D printing allows labs to customize their equipment to meet their own needs while reducing the cost of the equipment substantially compared with commercial systems.

It is possible to 3D print the entire housing for the imaging system as some researchers did for their systems (10,11,14,26–28), but we chose to use acrylic sheets and glue them together. Acrylic sheets are economical, ubiquitous, and easy to modify and can be assembled using acrylic plastic cement which allowed us to prototype several configurations and sizes before finalizing the implementation. Using glue enables fabrication of a significantly larger or small system without many changes by using larger or smaller acrylic sheets and allows for quick replacement of the imaging box sides if a different shelf configuration is needed. In our imaging box, all imaging components are attached to the lid making it easy to adjust the size parameters of the rest of the box.

### Control Electronics

The main components of the electronics are a Raspberry Pi 4, a logic level converter, eight NeoPixel RGB digital LED modules, and a Arducam IMX519 color camera (Table 1 in S1 for a complete list of components, suppliers, and costs). The logic level converter was used to convert the 3.3 V output from the Raspberry Pi to the required 5.0 V for the LED modules with an external power supply providing the necessary current, 0.5 A per LED at full power, for the LED modules. The NeoPixel LED modules included an integrated LED driver that allowed the LED modules to be connected in series and controlled by a pulse width modulated (PWM) output from the Raspberry Pi (Fig 8 in S3). The rated maximum power of each LED module was 3 W. The excitation wavelengths were measured using an Avantes AvaSpec-2048L Spectrometer with the peaks of the three different excitation wavelengths at 460 nm, 520 nm, and 630 nm (Fig 9 in S4). The control software on the Raspberry Pi microcontroller was developed in Python and used open-source libraries from Arducam and Adafurit to control the camera and LED modules (See S5 and S6 for installation and operation steps).

### Imaging and Analysis Software

Using FOSH and FOSH enables researchers to leverage open-source software libraries to control the equipment and image processing software to capture and process images for scientific analysis. We elected to use Python to control the camera and LEDs and to process the images due to the proliferation of application programming interfaces (API) and libraries available on the Raspberry Pi. Python is a powerful programming language that leverages the open-source community to develop and publish analysis software. Several research groups have published Python frameworks for computation biology, bio- and chemo-informatics, and fluorescence analysis (29–33). Our software leverages the published APIs for powering and changing the LEDs and capturing images from the camera. One choice open to the researcher is the image format for acquiring the image. The open source libcamera software on the Raspberry Pi allows the user to choose from several different image formats each with their own image processing levels. The formats available to the user are JPG, BMP, PNG, RGB, YUV, DNG and RAW. We focused on using PNG, DNG, and RAW file formats. PNG since it offered lossless compression with the built-in color balance, brightness, contrast, saturation, and bayer demosaic image processing already completed. The DNG format provides the use access to the raw image data with only bayer demosaic image processing performed. This allows the user to perform their own image processing for color balancing and level adjustment. The RAW file format offers the user access to the raw bit data from the camera sensor. This allows the user the most flexibility in processing the sensor data, but at the cost of complexity and file size.

There are multiple image analysis software tools available to researchers such as ImageJ, MATLAB, and Python to analyze. We chose to use Python for the image processing and analysis of the various experiments due to the open-source nature of Python. A benefit of using Python for the control software and the image processing software is that they can integrated into a single software application on the Raspberry Pi streamlining the image acquisition and result analysis allowing for real-time analysis of fluorescence results without a need for post experiment analysis.

### Imaging System Calibration and Performance

The color camera was tested and calibrated using two common test charts (Fig 2). A color calibration chart with reference colors and color codes, Fig 2A, was used to verify accurate color measurements which showed our system had close agreement between the published color values and the color data captured with the camera under natural light. We also tested response with a lens test chart that contained multiple bar, wedge, zone, and star patterns for lens distortion and resolution testing. Fig 2B shows that our camera and lens does not exhibit any barrel or pincushion distortion during image acquisition. Black levels of the camera sensor were tested by capturing a series of images inside the closed imaging box without any excitation light. The mean black levels across the full resolution of the camera in series of image was found to be [0.50, 0.36, 0.57] in RGB values with a standard deviation of [0.84, 0.68, 0.87] in RGB values. This indicated that no additional black level calibration was needed for the camera response. Fig 2C-F shows the effect of the different LED colors on the color response of the camera. Fig 2C was captured using all three excitation wavelengths at full power to generate a pseudo white LED color which caused a shift in the color balance levels of the camera since the wavelength levels were not balanced to produce balanced light. When using the blue, green, or red LEDs colors individually, Fig 2D-F, the color response of the image is influenced by the excitation wavelength used causing an overall shift in the color values in the image. The drive levels of each of the three difference LEDs were not balanced in power level which induced the shift in color levels measured from the color chart. When the imaging box is used for fluorescent images, optical filters will be used to separate the emission response from the excitation source and the shift in color balance from the camera does not influence the measurements. When the imaging box is used for full color scientific imaging the user will need to account for the color balance shift in their image processing pipeline to ensure that any measurements are not affected by the LED excitation (34).

The resolution of the imaging system was measured using a positive fluorescent 1951 USAF target, shown in Fig 3. The resolution was calculated at greater than 7-line pairs/mm with a line resolution at 70 μm with the diffraction limit of the camera at 2.3 μm. A line profile through group number 2 of the USAF target is available in the SI (Fig 10 in S4).

The linearity of the NeoPixel LEDs were measured using a ThorLabs PM13D optical power meter. The power meter was placed at varying distances below one of the LEDs in the imaging box and the LED brightness level adjusted from 0 to 100% in 10% increments. Fig 4A shows the measured power level for each of the excitation wavelengths with the 460 nm excitation showing the highest level of measured power and 620 nm showing the lowest level of measured power. All three excitation wavelengths exhibited a high level of linearity versus drive level with slight deviations at the at the 60-70% drive level for the 620 nm LED. Fig 4B shows the measure power in irradiance (W/cm^2^) as a function of distance from the LED. To determine the effect of continuous LED excitation on samples inside the imaging box a temperature logger was used to measure the change in ambient temperature in the box over the course of two hours. Fig 11 in S4 shows that the imaging box experienced an increase of 4 °F in ambient temperature when the LEDs were continuosly operated at maximum power for an extended duration. For experiments with longer excitation duration, care must be taken to ensure that the increase in temperature will not cause any damage to biological samples or turn off the LEDs between images.

**Fig 11.**
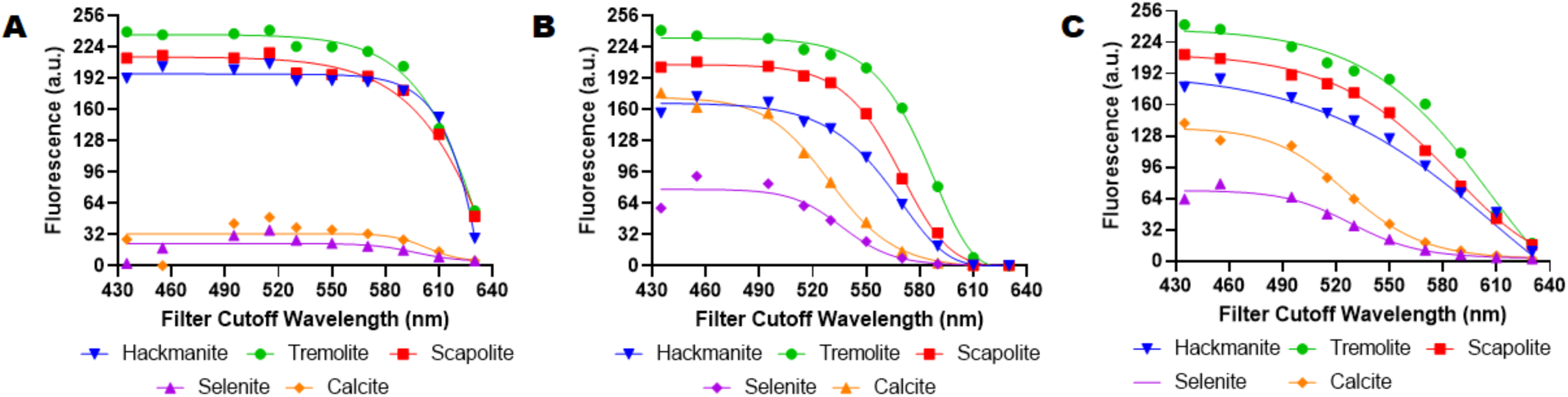
Fluorescence response of various minerals versus wavelength. (A) Fluorescence response in red channel (B) Fluorescence response in green channel (C) Monochrome fluorescence response

### Example Applications

#### Well Plate Imaging

The fluorescence sensitivity of the imaging system was tested using 200 µL serial dilutions from 10^-3^ M and 10^-12^ M of FITC in a 96-well PCR plate (Fig 5A). The imaging system was used to excite the FITC solutions with the 460 nm LED with an optical bandpass filter at 542 nm ± 25 nm (BrightLine) before the camera. The shutter time of the camera was varied from 1 ms to 10000 ms to determine the optimum exposure time to maximize the sensitivity measurements without overlading the camera sensor. The intensity values were extracted using Python and the data was analyzed using GraphPad Prism which showed that the imaging system could detect FITC concentrations from 1 mM to 10 pM (Fig 5D).

To test the fluorescence emission scanning capabilities of our imaging system sodium-sensitive (Fig 5B) and lithium-sensitive (Fig 5C) nanosensors were fabricated and characterized using serial dilution solutions of sodium chloride and lithium chloride respectively. The nanosensors were first characterized using a BioTeck Synergy H1 plate reader to get reference data using excitation and emission wavelengths that matched the LED excitation wavelengths and filter emission cutoff wavelength. The nanosensors were then imaged using our imaging system to determine the fluorescence emission capabilities of our system. The shutter time was varied from 10 ms to 200 ms based on the results from the FITC sensitivity measurements. As shown in Fig 5E and Fig 5D the imaging system performance matched the plate reader measurements. When dealing with a given concentration of fluorescent dye or a given dye brightness it is necessary to adjust the camera’s exposure time to maximize the measured dynamic range in the fluorescent response. For instance, a longer shutter time captures more light enhancing low dye concentration areas but sacrificing details in higher concentration areas. Similarly, a shorter shutter times can tune the dynamic range to focus on higher concentration responses by sacrificing performance in low dye concentration areas. This can be seen in Fig 5D where the exposure times influence the dynamic response of the various concentrations. For example, if a researcher is focused on FITC dye concentrations between 10^-3^ and 10^-6^ M an exposure time of 10 ms would give the best dynamic range for those concentrations. If a researcher was more interested in studying lower concentrations in 10^-8^ and 10^-12^ M range, an exposure time of 1000-2000 ms would tune the dynamic range to focus on the lower concentration values. In essence, adjusting the camera’s exposure time to the desired concentration or brightness range of the dye or subject would enable the researcher to achieve the best dynamic range in the image while maintaining consistency across experiments. These results indicates that our imaging system can be used to measure the fluorescence emissions of standard fluorescent dyes and analyte-sensitive nanosensors.

The flatness response of the imaging system was tested using 10 µM of FITC in each well of a 96-well plate (Fig 6A). The imaging system was used to excite the FITC solutions with the 460 nm LED with an optical bandpass filter at 542 nm ± 25 nm (BrightLine) before the camera. The shutter time was set to 10 ms based upon the results from the sensitivity tests (Fig 5). The intensity values were extracted using Python and the data was analyzed using GraphPad Prism which showed that the pixel value of fluorescence response only varied ±3% (Fig 6B) across the entire well plate.

#### DNA Electrophoresis Imaging

To determine the efficacy using openIVIS for DNA analysis, DNA gel electrophoresis was performed using a 1 kb DNA ladder with 10 µg in lanes 1-2 and 20 µg in lanes 3-4 (Fig 7). The completed gel was imaged using a Bio-Rad ChemiDoc Imaging system (Fig 7A) and our imaging system (Fig 7B) using a blue LED with a long pass 530 nm filter. Fig 7C shows a line profile through the left 10 µg lane in each image. Our imaging system was able to detect the different DNA base pairs in the agarose gel with similar performance to the commercial imaging system.

#### Animal Imaging

*In vivo* optical imaging of biological processes and phenomena is a powerful tool for researchers and educators. Leveraging optical imaging with animals or insects that have a short gestation and growth period allows researchers to study an animal’s lifecycle in a short period of time. Hornworms are the larvae of moths belonging to the family *Sphingidae* are commonly used as a model organism due to their large size, short life cycle, and simple physiology (35).

Tobacco hornworms undergo several stages of development, starting as eggs and progressing through several larval stages before pupating and eventually emerging as moths. During their larval stages, hornworms can grow quite large, with some species reaching lengths of up to 4 inches (36,37). Time-lapse imaging can provide high-resolution images of the animals over time to enable researchers to study various aspects of hornworm development, such as changes in body shape, size, and coloration (38,39). These images can provide valuable insights into the morphology and physiology of the hornworms and contribute to our understanding of their biology and ecology. To determine the feasibility of using our openIVIS system for time-lapsed imaging of a hornworm growth cycle we placed several hornworms in our imaging box and took pictures every thirty minutes for fourteen days. Fig 8 shows six of the images taken over the experiment timeline to highlight the ability of our imaging system to perform time-lapsed monitoring of a hornworm’s growth cycle. The minimum imaging period is a combination of user settings such as preview options, image resolution, and Raspberry Pi storage write speed with typical still image captures times on the order of several seconds. It is possible to export the camera outputs as video frames with video rates up to 100 frames per second (FPS) at lower resolutions enabling time lapsed images at 10 ms intervals.

#### Plant Health Imaging

Fluorescence imaging can be used to measure several parameters related to plant physiology such as environmental stress, photosynthetic efficiency, nutrient deficiency, and disease (40–43). By measuring these parameters, researchers and agriculturists can gain insights into the health and performance of the plant under different environmental conditions where early detection of diseases and pests can save crops and improve yield. When plants are under stress, the fluorescence response of different pigments and compounds present in the plant tissue, such as chlorophyll, anthocyanins, and carotenoids change across different plant tissues (40–43). Anthocyanins are pigments that give plants, including lettuce, a red, purple, or blue color and protect plants from harmful radiation. Using the anthocyanin index formula (40) we calculated the anthocyanin response of a lettuce imaged over several days in our openIVIS system (Fig 9). As expected, the anthocyanin index clearly changes over time as the lettuce leaf becomes dehydrated in the imaging system.

#### Mineral Spectral Response Imaging

Mineral fluorescence is the phenomenon of a mineral emitting light when exposed to certain types of radiation, such as ultraviolet (UV) light. This fluorescence can be used to detect and identify minerals, as each mineral has a unique spectral response when exposed to UV light (44,45). Mineralogists often use spectrometers to measure the spectral response of minerals. To demonstrate the feasibility of using our openIVIS system to measure the spectral response of select minerals we replaced the NeoPixel RGB LEDs with 10 W long wave UV (395 nm) LEDs (Amazon, Part Number: B0863KSRKB) and used manual power control rather than the electronic control from the Raspberry Pi of the light source. We imaged nine minerals (Amazon, Part Number: B00FGDOW1O) with eleven long pass optical filters and measure the fluorescence response of the minerals. The RGB channels for each mineral were analyzed for the fluorescence response in each channel. In addition, the RGB data were converted to grayscale using a Python OpenCV library to look at a combined fluorescence response. The minerals shown in Fig 10 are as follows (1) Hackmanite from Canada, (2) Scapolite from Canada, (4) Calcite from Montana, (7) Selenite from Texas, (10) Fluorite from Utah, (11) Turritella Agate from Wyoming, (13) Tremolite (unspecified source), and (14) Calcite from Mexico. Comparing the spectral response (Fig 11) of hackmanite, tremolite, and scapolite to selenite and calcite shows it is possible to image minerals based on their spectral response using our openIVIS system.

### Potential openIVIS Modifications

There are several modifications that we have considered for future investigations or modifications that other groups could perform with their own version of openIVIS.

#### Raspberry Pi Alternatives

While the Raspberry Pi series are the best known and most widely used small single board computers (SBC) due to their versatility, reliability, ease of use, low power, and price there are many alternatives that researchers can use for their IVIS systems each with its own strengths and weaknesses. We selected a Raspberry Pi for our implementation since it was the standard approach from other similar projects. Other researchers might have other needs for their SBC to expand their openIVIS systems or have challenges in acquiring a Raspberry Pi, and in the SI (S2) we detail several SBCs that are options for consideration.

#### Camera Selection

The scientific community is embracing FOSS and FOSH concepts to enable the proliferation of laboratory equipment with hundreds of researchers designing and making their own equipment. The use of open-source hardware allows researchers the ability to freely swap out individual pieces of the equipment dependent on the performance level needed for the experiment. The open-source nature of our imaging system aims to leverage existing hardware and software from the non-traditional scientific community to reduce the hardware and software development time. The use of FOSH in the imaging system allows the user to choose from several different camera types based upon the intended application. The Raspberry Pi has built in libraries for handling both monochrome and color cameras with options from 1 megapixel (MP) to 64 MP depending upon the need. Color cameras use a color filter array (CFA) above the sensor array to provide red, blue, and green color information using bandpass filters for each color range. Monochrome cameras do not have a CFA above the sensor array with each pixel seeing the entire wavelength spectrum. Since monochrome cameras do not have a filter, they provide better performance in low light conditions since all the light reaches the sensor.

Color cameras offer higher resolutions and lower cost due to the wide use of color cameras in normal situations. Color cameras allow the user to look at different wavelength responses simultaneously which enables tracking different fluorescent and stain responses at the same time. The choice of monochrome versus color depends upon the intended application. We chose to use a color camera, Arducam IMX519, due to the increased flexibility in looking at both fluorescent samples and live animals. The Arducam IMX519 color camera is a 16 MP autofocus high-resolution camera module. The autofocus lens on the camera allowed the imaging system to automatically adjust camera focus on samples placed at different heights that allowed the users to maximize the imaging area of the objects for each experiment. Some cameras enable the user to swap out lenses to change the field of view (FOV) of the camera enabling the user to select the desired imaging area and magnification for their specific application (S7).

Several other factors to consider when choosing a camera is sensitivity, resolution, and frames per second which have an influence on each other. The sensitivity of a camera is typically measured in terms of its quantum efficiency, which is a measure of the percentage of photons that the camera can convert into electronic signals. High quantum efficiency means that the camera will be able to capture more photons, resulting in higher signal-to-noise ratios and better image quality in low light conditions. The resolution of a camera refers to the number of pixels it has, which determines the level of detail that it can capture. Frames per second, on the other hand, refer to the number of frames that the camera can capture in a given amount of time. The maximum number of frames per second that a camera can capture will depend on several factors, including the camera’s sensor size, readout speed, and exposure time. Some cameras are capable of capturing tens or even hundreds of frames per second, while others may only be able to capture a few frames per second.

Higher resolution cameras can capture more detail in an image, but they may also have smaller pixels, which can reduce sensitivity. This is because smaller pixels can capture fewer photons, which can result in lower signal-to-noise ratios and reduced image quality in low light conditions. However, higher sensitivity cameras may also have larger pixels, which can reduce the overall resolution of the image. Higher resolution cameras can capture more detail in an image, but they may also require more time to process the data, resulting in slower frame rates. In general, higher resolution cameras tend to have slower frame rates, while cameras with lower resolutions can achieve faster frame rates.

When selecting a camera, it is important to consider the specific requirements of your experiment and choose a camera that has the appropriate sensitivity, resolution, and maximum frames per second for your needs. In general, the best camera for a particular application will depend on the specific requirements of that application. For example, if the application requires high-resolution images of a object with high fluorescence, a high-resolution camera with smaller pixels may be the best choice. If the application requires high-quality images in low light experiments, a more sensitive camera with larger pixels may be the better choice, even if it has a lower overall resolution. If an experiment requires slow or static fluorescence responses, a higher resolution camera with a lower frame rate may be sufficient. However, if the application requires capturing fast fluorescence motion to be captured with high temporal detail, a camera with a lower resolution but a higher frame rate may be the better choice.

Another potential modification is the use of multiple synchronized cameras in the box that would allow for stereoscopic imaging. Arducam offers several multiplexer and stereo cameras for the Raspberry Pi SBCs. Incorporating a synchronized stereo camera system that uses two or more cameras precisely synchronized to capture simultaneous images of the same imaging object from slightly different perspectives could be combined to create a 3D representation of the object. Synchronization is critical for accurate 3D reconstruction, as any difference in timing between the cameras can result in misalignment of the images and errors in depth perception. The Arducam boards automatically synchronize the cameras enabling researchers to focus on the application versus the implementation of the stereoscopic imaging system.

#### Imaging Box Design

In the development of the Open IVIS system we implemented an approximately cubical box for simplicity and portability. The potential shapes and sizes of the imaging system housing is only limited by a researcher’s 3D printer or fabrication facility. The system housing could be made wider to accommodate bigger sample sizes, or smaller if a researcher did not need a big imaging area. The imaging box could be made from various materials such ranging from interlocking wood panels (Fig 13 in S8) to cardboard (Fig 14 in S8) to LEGO blocks (Fig 15 in S8, not recommended due to the price difference between LEGO and acrylic sheets). In the case of LEGO blocks, images can be captured using this type of setup (Fig 16 in S8), but a researcher needs to ensure that any gaps between blocks are sealed for experiments when low levels of light are being measured. The housing does not need to remain a rectangle, for example a dome shaped housing could be made that would still allow for equal LED excitation throughout the box. The tradeoffs in housing size are portability, weight, ease of fabrication, and flexibility in imaging sample size.

Another factor to consider when designing an imaging box is the type of excitation source needed and the type of fluorescent measurements that are needed(46). There are several other potential excitation light sources that can be used based on the desired measurements, including single-color LEDs, white light sources, and different arrays of LEDs. Single-color LEDs are commonly used in applications that require a specific wavelength of light for excitation, such as fluorescence images. These LEDs emit light at a specific wavelength, which can be chosen based on the fluorescent molecule being used. Single-color LEDs are also relatively inexpensive, consume less power, and have a long lifetime. The LEDs can be arranged in an array (11,13,47,48) or in single source configurations (10,40,49) based upon the measurements that are desired. If absorption measurements are needed the sample needs to be placed between the excitation source and the camera, while fluorescence emission measurements require the LEDs and camera to be on the same side of the object being imaged. LED arrays are typically seen in instruments that are used for absorption. With fluorescent measurements a diffuser can be place over the LEDs to reduce or prevent any light reflections as seen in Fig 8 that may disrupt readings. Different arrays of LEDs, such as multichannel LED arrays, can be used to provide excitation at multiple wavelengths simultaneously. These arrays can be customized to provide specific wavelengths of light, making them useful in a wide range of applications, including fluorescence microscopy, flow cytometry, and medical imaging. White light sources, on the other hand, emit light across a broad range of wavelengths, which can be useful for applications that require excitation across multiple wavelengths and when optical filters can be used to provide wavelength selectivity (15,50). White light sources can be produced using different technologies, including halogen lamps, xenon lamps, and LEDs. However, the use of white light sources for excitation can be limited due to their relatively low intensity and lack of spectral specificity. Overall, the choice of excitation light source depends on the specific application and the requirements for wavelength, intensity, and spectral specificity. Single-color LEDs are a good choice for applications that require specific wavelengths of light, while white light sources and different arrays of LEDs can be used for applications that require excitation across multiple wavelengths.

## Conclusion

In this study we designed and manufactured a triple-excitation, multi-emission open-source imaging system that can be quickly adapted for laboratory experiments studying fluorescence in biological models. We demonstrated the capabilities of this system while exploring the fluorescence response of several ion selective nanosensors, a DNA ladder in an agarose gel, various mineral spectral responses, plant health, and live animal time-laps imaging. The system utilized an integrated three-color LED matrix and a commercialized autofocus high-resolution color camera. While the spectral range is limited to the wavelengths of the LED elements in the LED matrix, the flexibility of quickly interchangeable optical filters enables the user to measure a wide range of fluorescent responses in various experimental setups. The spectral sensitivity of our system able to measure nano to picomolar concentrations of various analytes. The flexibility of the Raspberry Pi framework allows the user to easily swap out cameras and lenses to increase or decrease the imaging region and spatial resolution. The Raspberry Pi general purpose input output (GPIO) pins enable a user to choose from a wide range of excitation sources from LEDs to lasers. The open-source nature of the imaging system enables interdisciplinary collaboration across multiple levels of STEM education and research. Importantly, our system takes a step in enabling fluorescent imaging in a wide range of environments from the laboratory to the high school classroom.

## Supporting information

Open IVIS Supplementary Information

## Acknowledgements

The authors thank Susanta Sarkar in the Physics department at the Colorado School of Mines for the use of their DNA gel imager and Anneke Cash for assistance in building the LEGO imager. The authors thank Tyler Sodia and Adrian Mendonsa for their careful reading of the manuscript.

