## Supplementary material for "Multifunction Fluorescence Open-Source *In Vivo/In Vitro* Imaging System (openIVIS)": Open IVIS Supplementary Information

[2] The MITRE Corporation, Bedford, Massachusetts, United States of America. The author's affiliation with The MITRE Corporation is for identification purposes only and is not intended to convey or imply MITRE's concurrence with, or support for, the positions, opinions, or viewpoints expressed by the author. Approved for Public Release, Distribution Unlimited. Public Release Case Number 23-1621.

[3] Chemical and Biological Engineering Department, Colorado School of Mines, Colorado, United States of America

[3] Chemical and Biological Engineering Department, Colorado School of Mines, Colorado, United States of America

[4] Mechanical Engineering, Colorado School of Mines, Colorado, United States of America

[5] Electrical Engineering, Colorado School of Mines, Colorado, United States of America

[6] Arvada West High School, Arvada, Colorado, United States of America

[7] Colorado Early Colleges Castle Rock, Castle Rock, Colorado, United States of America

[8] Department of Engineering, Harvey Mudd College, Claremont, California, United States of America

### S1. Imaging Box Information

openIVIS CAD files and STL files are available on GitHub at the following link:

https://github.com/CashLabMines/openIVIS.git

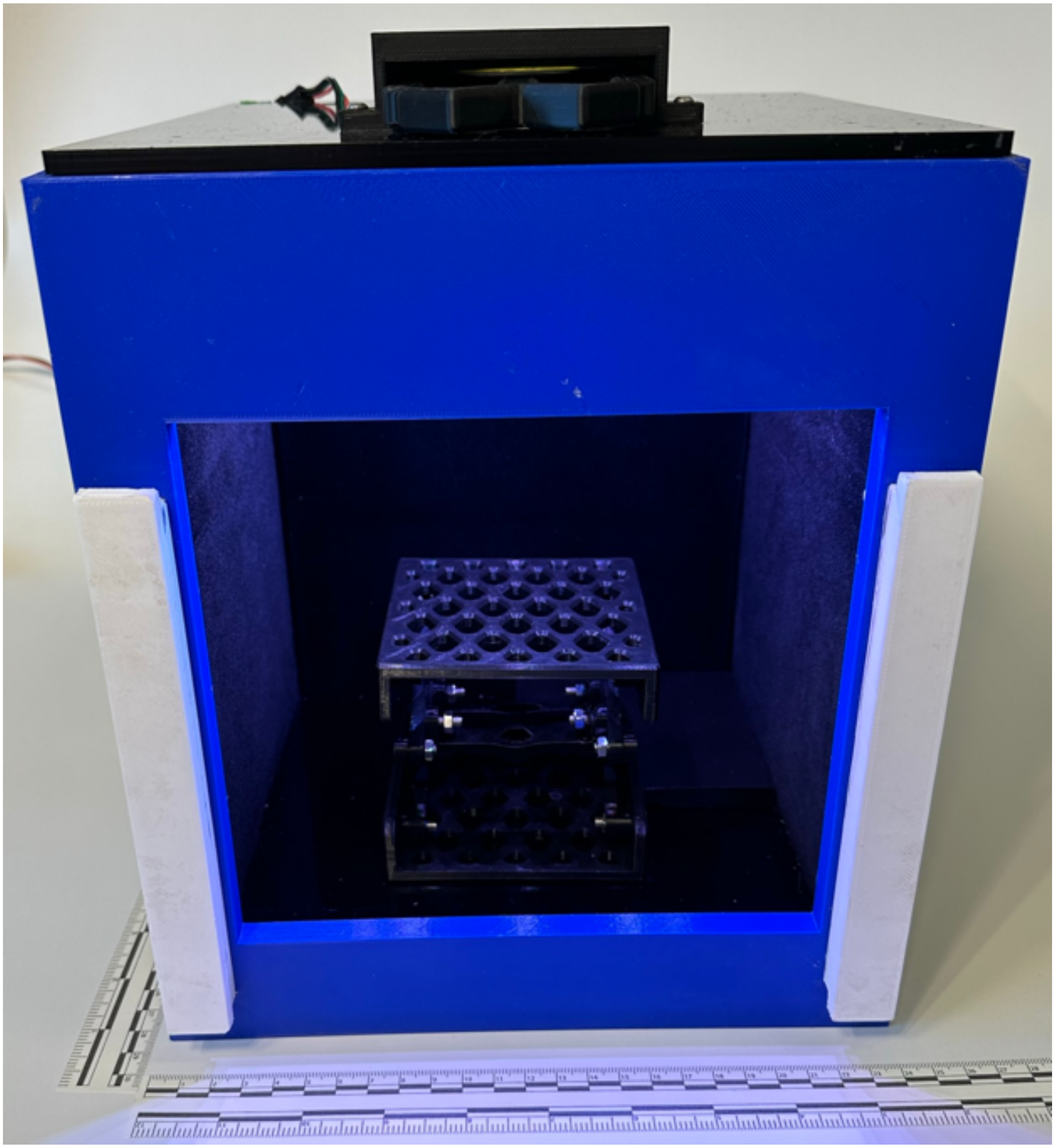

**SFig 1 openIVIS Interior View**

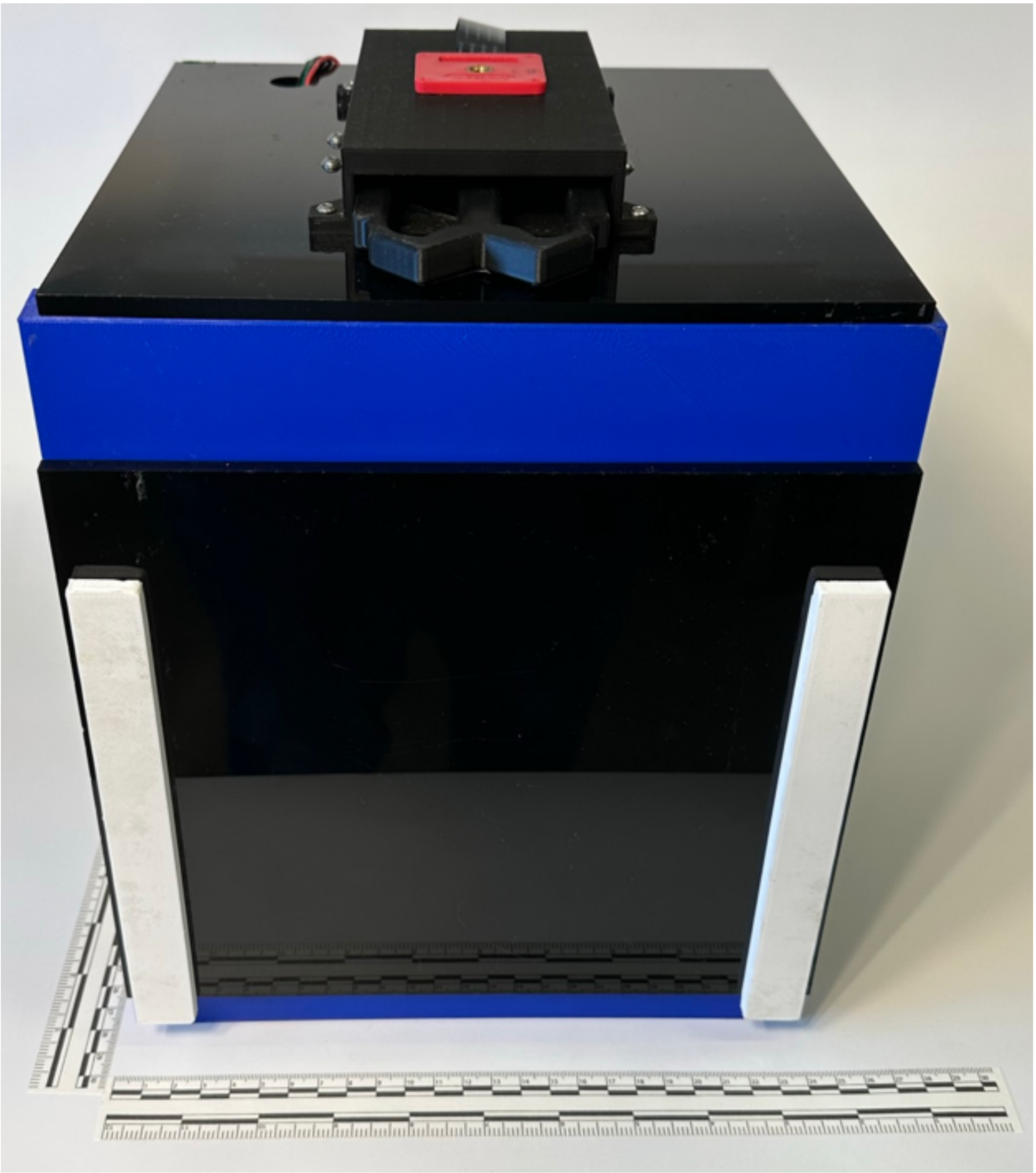

**SFig 2 openIVIS Front View**

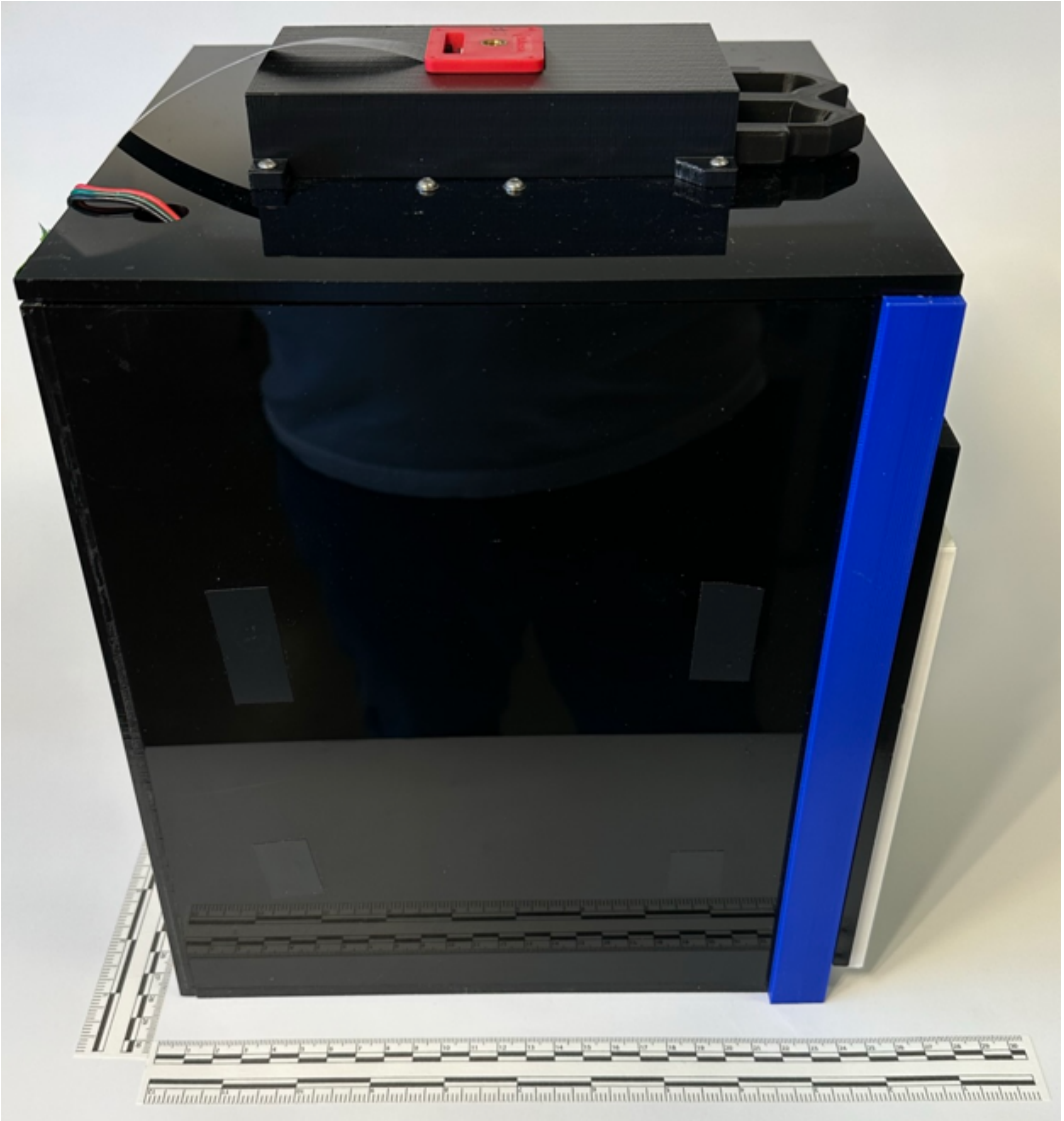

**SFig 3 openIVIS Lateral View**

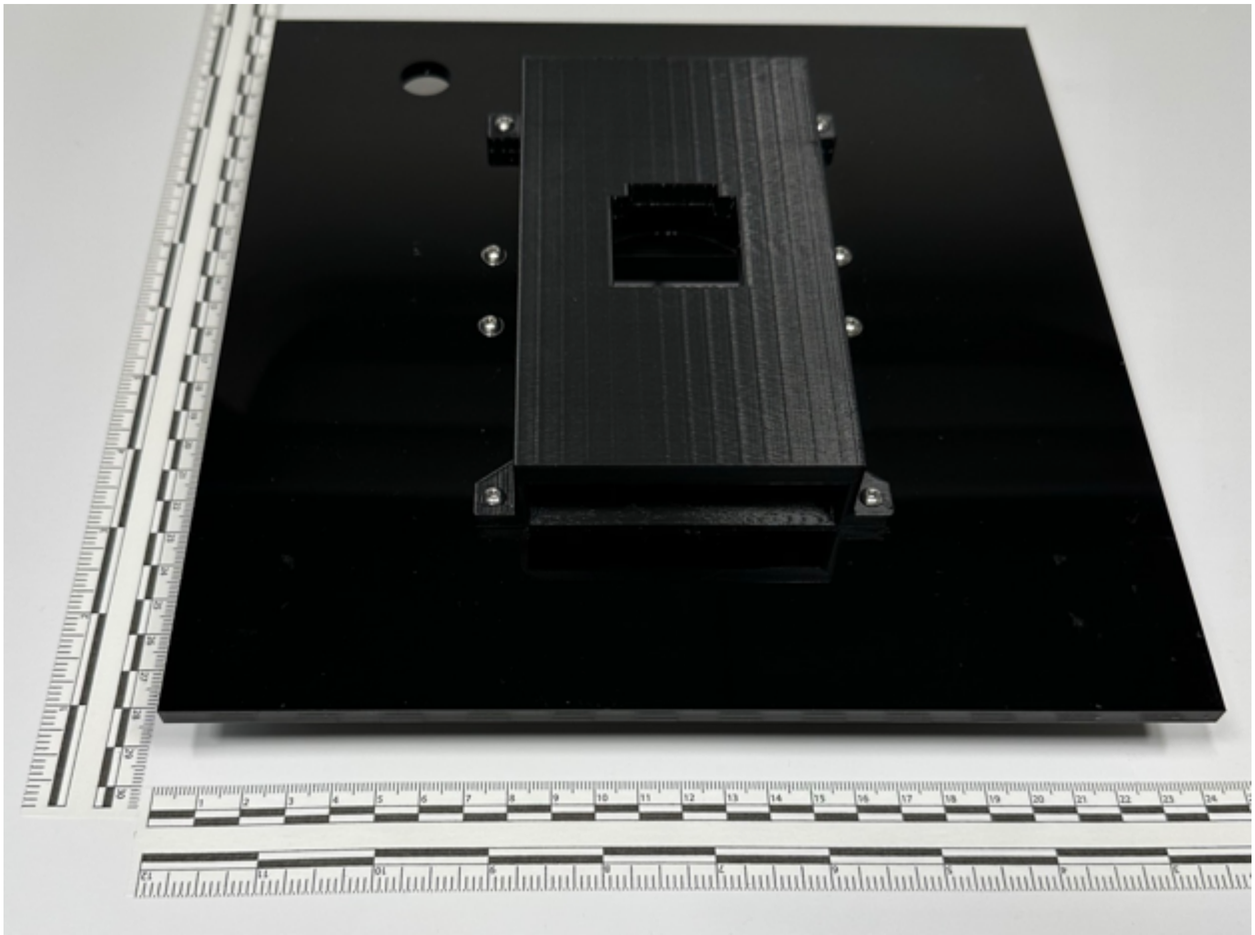

**SFig 4 openIVIS Camera Holder**

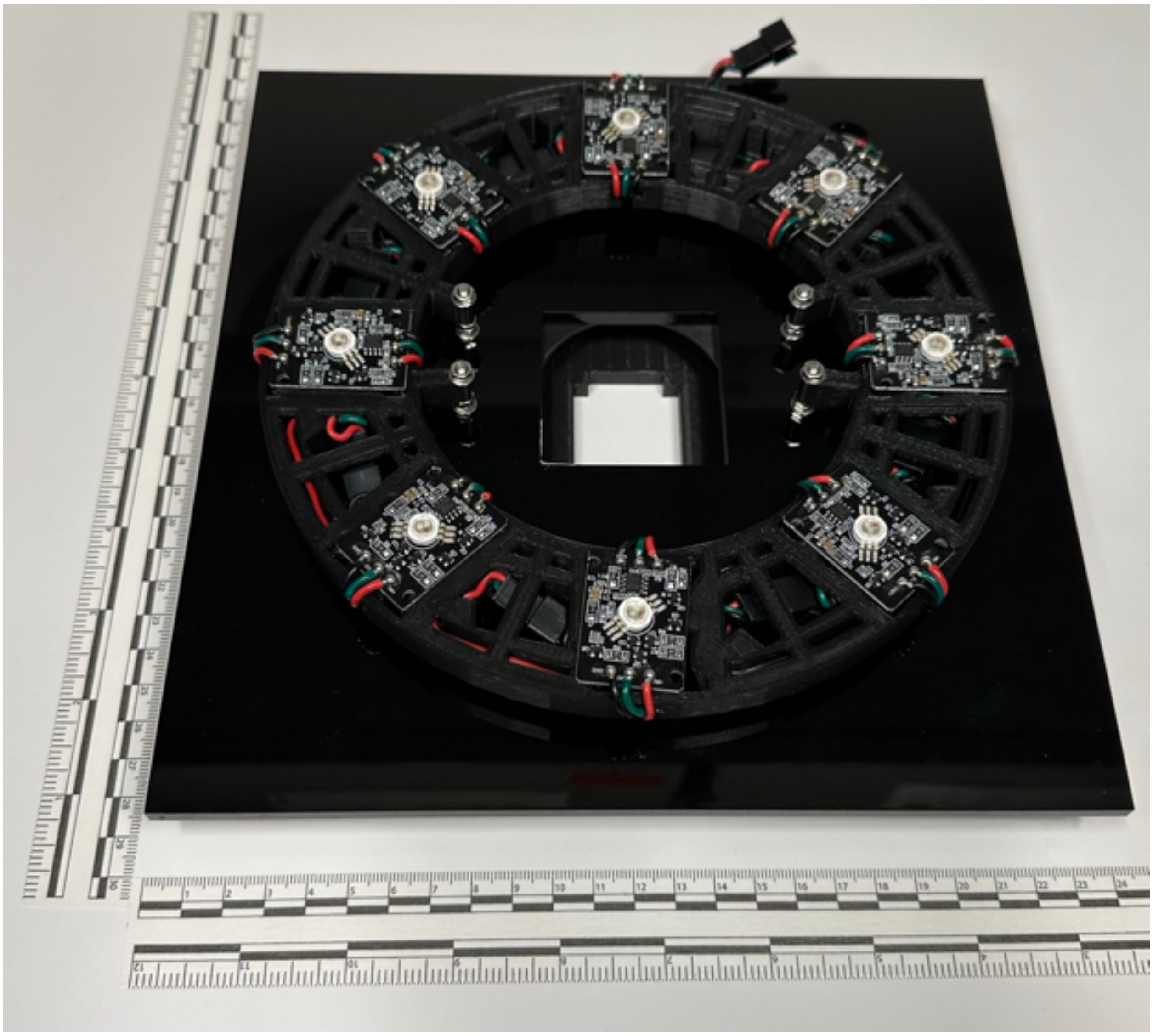

**SFig 5 openIVIS LED Holder**

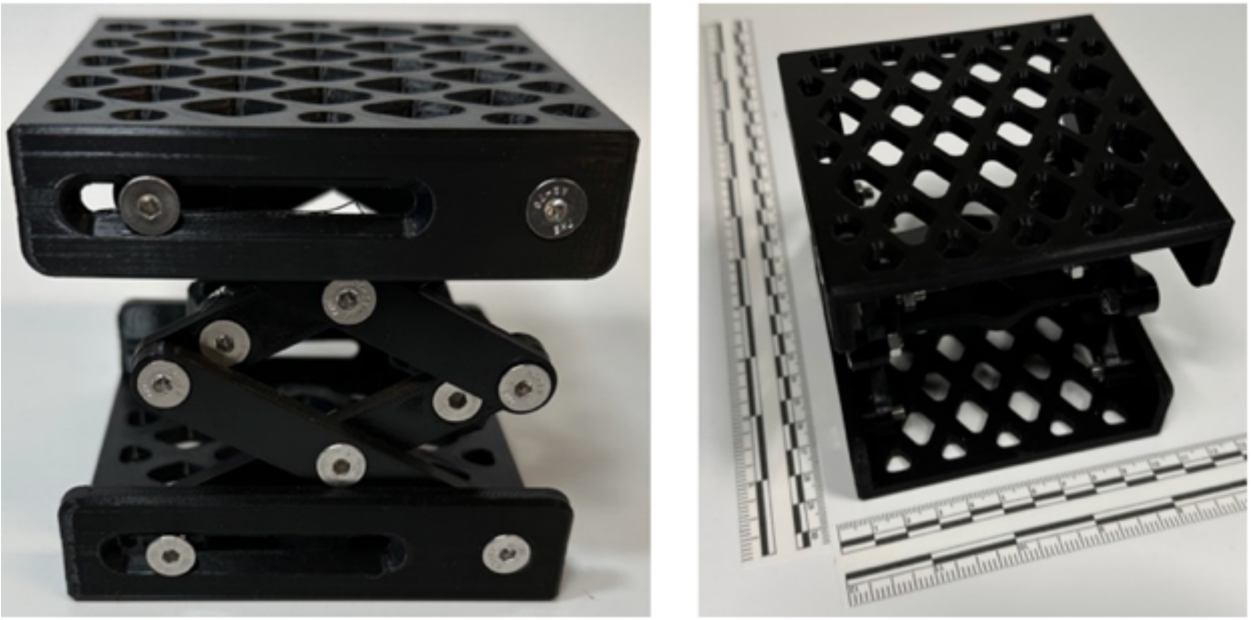

**SFig 6 openIVIS Adjustable Stand**

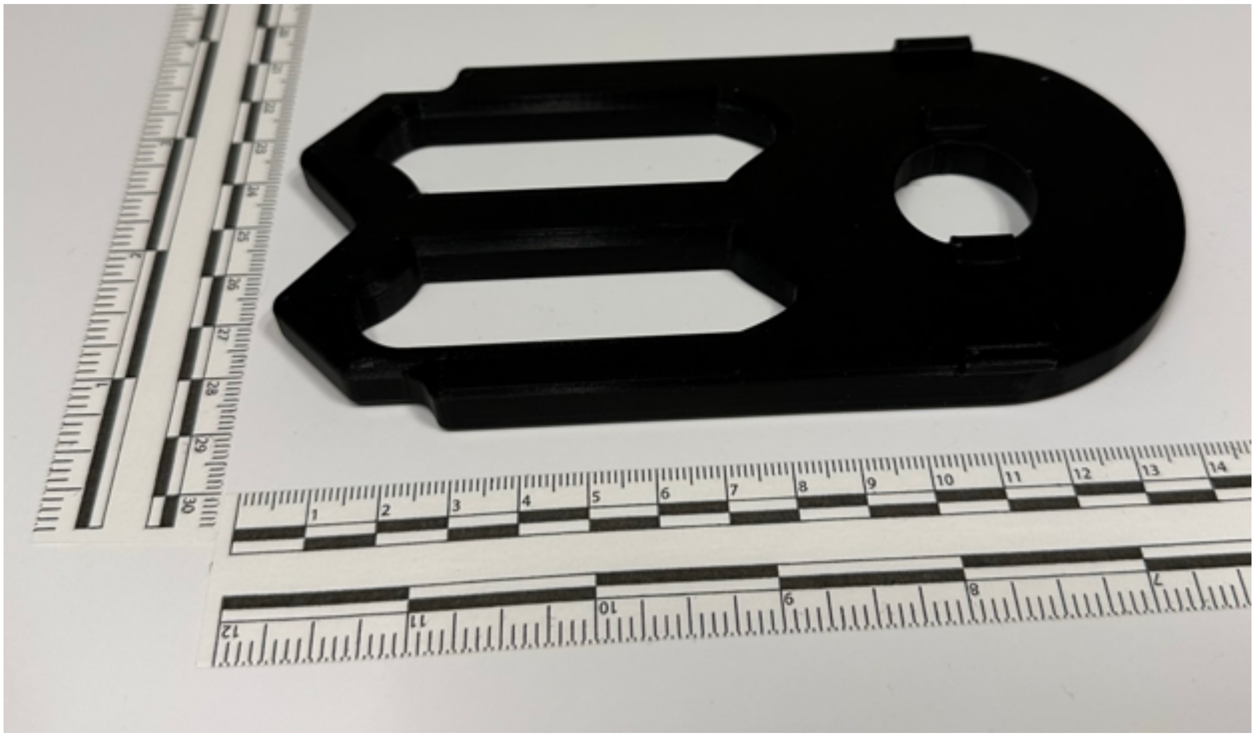

**SFig 7 openIVIS Filter Holder**

**STable 1 Open IVIS Bill of Materials**

| Component | Supplier | Part Number | Unit Cost | Qty | Total Cost |
|---|---|---|---|---|---|
| Raspberry Pi 4, 4 GB Ram | CanaKit | PI4-4GB-STR32F-C4-BLK-1 | $119.95 | 1 | $119.95 |

| | | | | | |
|---|---|---|---|---|---|
| Samsung Pro Plus Micro SSD, 256 GB | Amazon | MB-MD256KA/AM | $20.88 | 1 | $20.88 |
| Arducam IMX519 Camera | Arducam | B0371 | $24.99 | 1 | $24.99 |
| Ardufruit Camera Flex Cable, 24" | Adafruit | 1731 | $2.95 | 1 | $2.95 |
| NeoPixel LED, RGBW, 4W, Cool White | Adafruit | 5162 | $5.50 | 8 | $44.00 |
| Raspberry Pi Prototype Board | Adafruit | 2314 | $7.50 | 1 | $7.50 |
| Logic Level Converter, BSS138 | Adafruit | 757 | $3.95 | 1 | $3.95 |
| 3-Pin Receptacle Cable | Adafruit | 1663 | $1.50 | 1 | $1.50 |
| 2.1 mm DC Barrel Jack | Adafruit | 373 | $0.95 | 1 | $0.95 |
| Stranded-Core Wire, Black, 22 AWG | Adafruit | 2976 | $2.95 | 1 | $2.95 |
| Stranded-Core Wire, Red | Adafruit | 3068 | $2.95 | 1 | $2.95 |
| Stranded-Core Wire, Yellow | Adafruit | 2987 | $2.95 | 1 | $2.95 |
| Brass M2.5 Standoffs for Pi HATs | Adafruit | 2336 | $0.75 | 2 | $1.50 |
| Adhesive Black Felt Fabric | Amazon | B08DS6CZRJ | $0.75 | 4 | $3.00 |
| Black Acrylic Sheets (12” x 16”) | Amazon | B0BCFVVMYW | $8.65 | 6 | $51.90 |
| Power Supply, 12V 5A | Amazon | B08C594VNP | $10.99 | 1 | $10.99 |
| | | | | **Total Cost** | **$302.91** |

### S2. Alternative Single Board Computers

Alternative single board computers to the Raspberry Pi are discussed below. These are just several examples of the many alternatives to the Raspberry Pi that are available to developers and researchers. Ultimately, the choice of which board to use depends on your specific needs and requirements.

- ASUS Tinker Board: This is another single-board computer that is like the Raspberry Pi. It has a faster processor and more RAM, making it a good option for running more demanding applications(1).
- Nvidia Jetson Nano: The Jetson Nano is a powerful single-board computer designed for AI applications. It has a fast CPU and GPU and can handle advanced AI workloads. It is more expensive than the Raspberry Pi but offers better performance for certain applications(2).
- Libre Computer Board AML-S905X-CC (Le Potato): This is another single-board computer that is much like the Raspberry Pi. It has a similar processor and RAM and comes with a GPU. It has similar peripheral support making it an good alternative to the Raspberry Pi(3).

- BeagleBone Black: Like the Raspberry Pi, the BeagleBone Black is a small, single-board computer that can run Linux and can run a variety of applications. It has more built-in connectivity options and a faster processor than the Raspberry Pi(4).
- Odroid XU4: This is a powerful alternative to the Raspberry Pi that has a faster processor and more RAM. It is great for running more intensive applications such as media servers, gaming, and emulation(5).

### S3. NeoPixel LED Driver Board Schematic

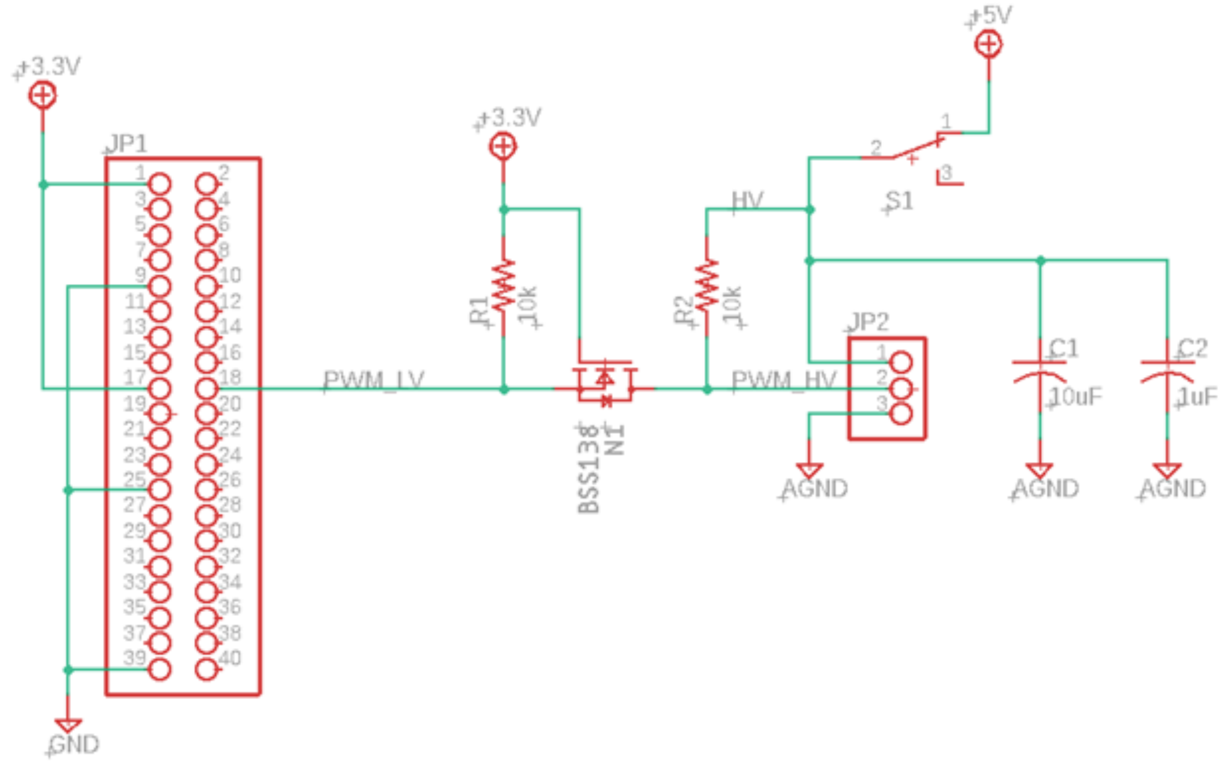


**SFig 8 NeoPixel LED driver board schematic**

### S4. Imaging Box NeoPixel LED Calibrations

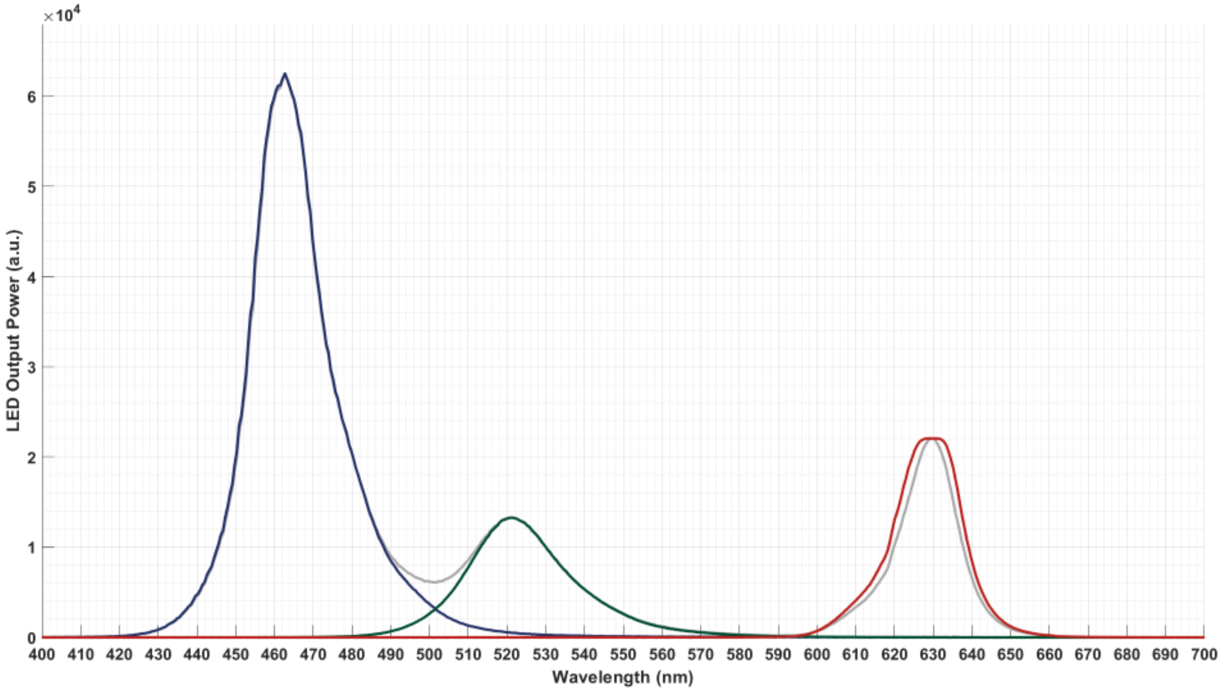


**SFig 9 NeoPixel LED Spectrum**

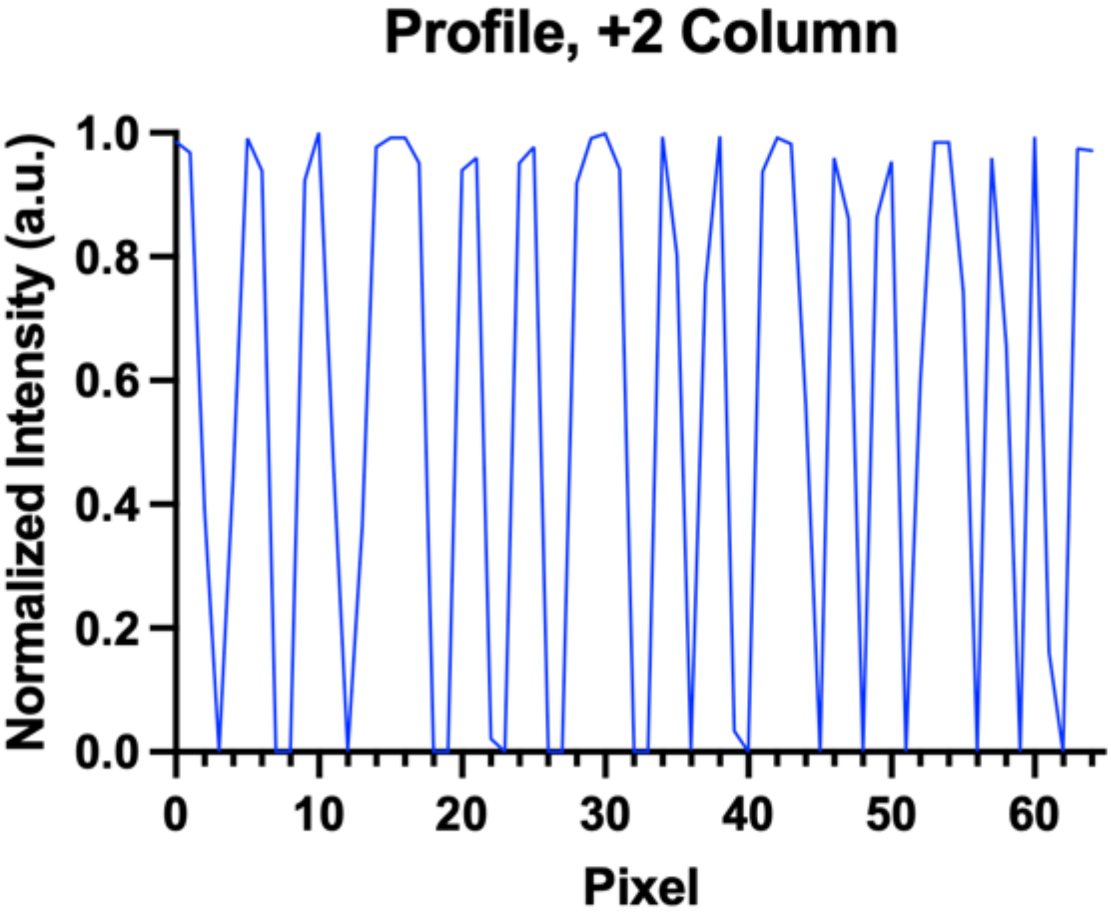


**SFig 10 USAF Target Line Profile**

To determine the effect of the LEDs on the temperature inside the imaging box during long excitation durations a temperature logger (Elitech GSP-6) was used to record the temperature in ten second intervals while each of the eight LEDs were set to cycle through random colors for two hours. The temperature in the box increased three degrees over the course of the two hours with the temperature continuing to increase when the recording was stopped.

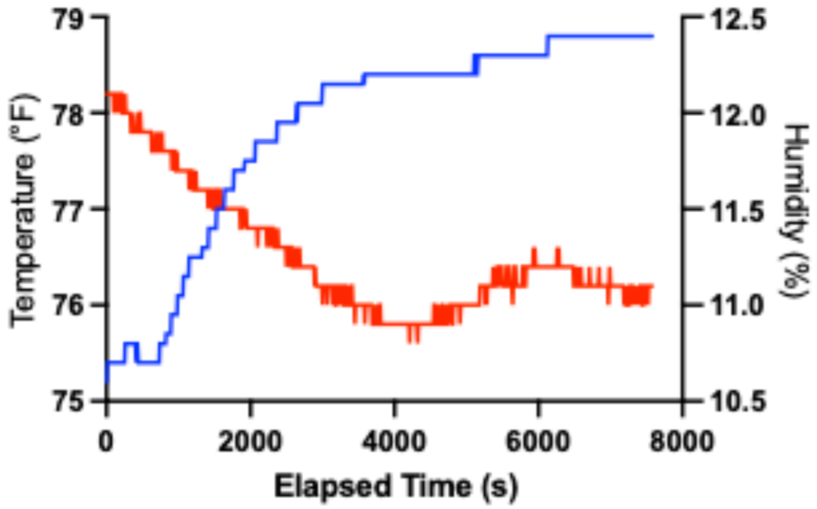


**SFig 11 Temperature and Humidity Response inside imaging box with all eight LEDs on**

### S5. Initial Configuration Steps

There are several installation steps needed to get the Arducam IMX519 drivers installed on a Raspberry Pi. Adrucam has a Getting Started guide on their website(6). This guide provides instructions on how to install the IMX519 camera drivers and how to enable them in the boot config file. Adafruit has a guide on their website that provides and overview with instructions on how to control NeoPixel LEDs with Python on a Raspberry Pi(7).

### S6. System Operation

On the Raspberry Pi the python code is stored in /home/pi/Python. The open a terminal window either select the terminal icon on the menu task bar or via the start menu through "Accessories > Terminal". Two methods to execute python scripts:

- In a terminal window type "cd /home/pi/Python", then to execute a python file type "sudo python <filename>" in a terminal window where <filename> is the full python file name such as "fluorescence.py"
- For the standard scripts listed below, the "sudo python <filename>" was added as an alias in the ".bashrc" file in the /home/pi/ directory. This lets the user execute the commands from any directory.

**Experiment Operation**

For Fig 2, Fig 3, and Fig 8 the camera was operated using the command libcamera-still as follows: libcamera-still --autofocus --vflip --hflip -n --immediate -e png -o [filename]

For Fig 5, Fig6, Fig 7, Fig 9, and Fig 10 the camera was operated using the command libcamera-still as follows: libcamera-still --autofocus --vflip --hflip -n --immediate --raw 1 --shutter [XXX] -o [filename] where the shutter value was looped through the values 1, 2, 4, 6, 8, 10, 20, 40, 60, 80, 100, 200, 400, 600, 800, 1000, 2000, 4000, 6000, 8000, 10000 to adjust the camera exposure time in milliseconds.

The LEDs were controlled using the NeoPixel library in python using the following commands:

```
pixels = neopixel.NeoPixel(board.D18, 8, brightness=1.0, auto_write=False,
pixel_order=neopixel.RGB)
pixels = (desired color)
pixels.brightness = 1.0
pixels.show()
```

The desired colors are controlled by a three-integer value variable to set the RGB color. For blue LED (460 nm) excitation: pixels = (0,0,255), for green LED (520 nm) excitation: pixels = (0,255,0), for red LED (630 nm) excitation: pixels = (255,0,0), for white LED excitation: pixels = (255,255,255), for external excitation such as UV: pixels = (0,0,0). To change the LED power level the pixels.brightness was changed to a value between 0 – 1.0 based upon the desired output power.

**Camera Operation**

There are two methods to open a video stream from the camera. Note the "--autofocus" command syntax is dependent on version of the libcamera library installed and will vary slightly. The command "libcamera --help" will provide the details on the command syntax.

- Alias name "camera". Type this command in any terminal window to execute the libcamera-script. This will open a video stream from the camera. To stop the video stream either close the video window or type "Ctrl+C" in the terminal window to terminate the command.
- Alias name "libcamera-still". Type this command in any terminal window to execute the libcamera-script. There are many options with the script and user can type "libcamera-still --help" in a terminal window to see the various options.
    - Preview window: "libcamera-still --autofocus --vflip --hflip -t 0" this will open a video stream without a timeout, perform autofocusing, and perform a vertical and horizontal flip of the image to align the image with the back of the box at the top of the image.
    - Save image: "libcamera-still --autofocus --vflip --hflip --e png -o <filename>" this will perform autofocusing, perform a vertical and horizontal flip of the image to

align the image with the back of the box at the top of the image, and save the image based on the user input <filename>

To edit python files, open the “File Manger” icon on the task bar. Change directory to “/home/pi/Python”. Right-click on the python file the user wants to edit and select “Geany”. This will open a built-in simplified IDE that allows the user to edit the python files.

**Standard Python files**

Python files with alias commands in the “*.bashrc*” file.

- **lightdimmer.py** : this one cycles through the LED colors with a 1-2 sec delay for each color. When the desired color is turned on the user needs to type “Ctrl+C” to cancel the python script. The LEDs will remain on with the last color. Alias name “dimmer”. Type this command in any terminal window to execute the python script.
- **fluorescence.py** : this one is the main imaging script for fluorescence measurements. The LED excitation color is controlled by the variable “colorLoop” on line 31 of the file. To add additional colors to the script, replace the number with the desired color, 0 = Red, 1 = Green, 2 = Blue, 3 = While, 4 = UV or external illumination. To execute multiple colors, the user needs to add a comma between colors, i.e. [0,1,2] to use Red, Green, and Blue excitation. The script will take both .png and .dng pictures for each exposure setting. The exposure settings are controlled by the variable *scale* on line 19 in the python file “saveImages.py”. The python script will ask the user for the experiment name. The experiment name will be used as the base string for each of the images with the LED color and exposure time appended to the image filename. Alias name “fluorescence”. Type this command in any terminal window to execute the python script. The python scripts save the images to “/home/pi/Pictures/<experiment name>” where <experiment name> is the user entered string when executing the python script.

### S7. Field of View Lenses

Images of the calibration chart taken with an Arducam OV2311 monochrome camera (UCTRONICS, Part Number B0381) with Arducam M12 Lens set (UCTRONICS, Part Number LK001). The camera was positioned 260 mm from the calibration chart for each picture.

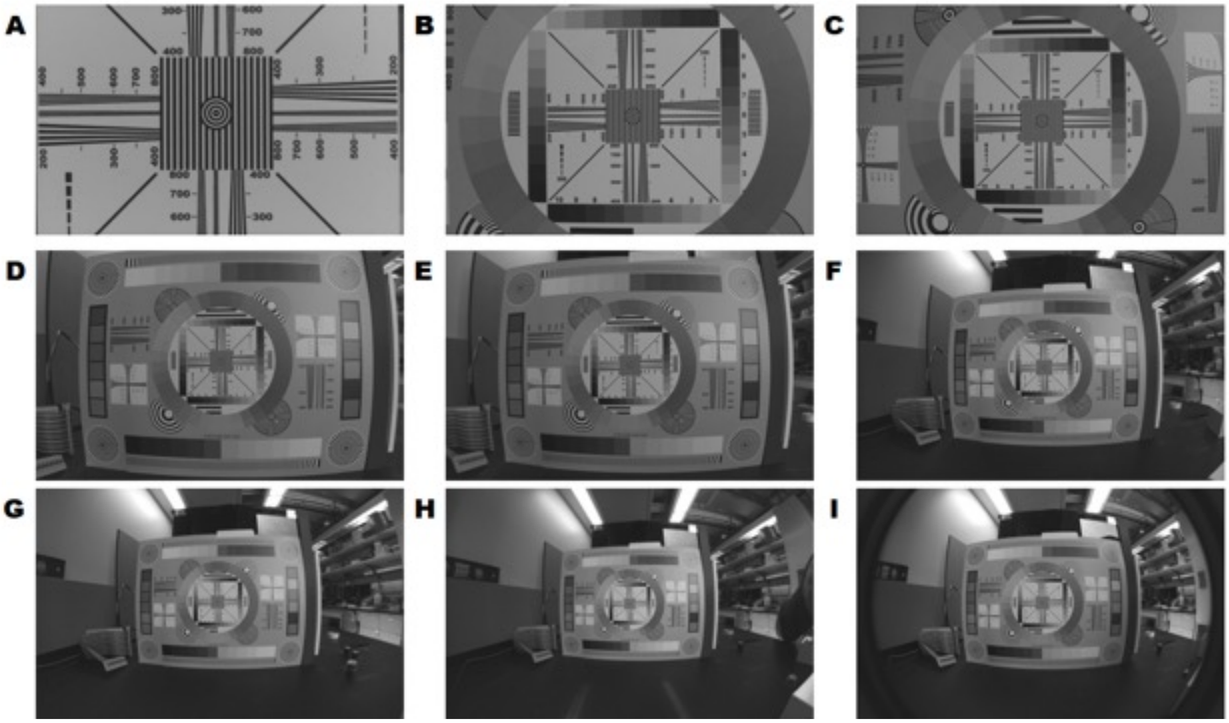


**SFig 12 Image of calibration chart at 260 mm with different field of view lenses (A) 10° FOV (B) 20° FOV (C) 40° FOV (D) 60° FOV (E) 80° FOV (F) 100° FOV (G) 120° FOV (H) 140° FOV (I) 160° FOV**

### S8. Alternative Box Designs

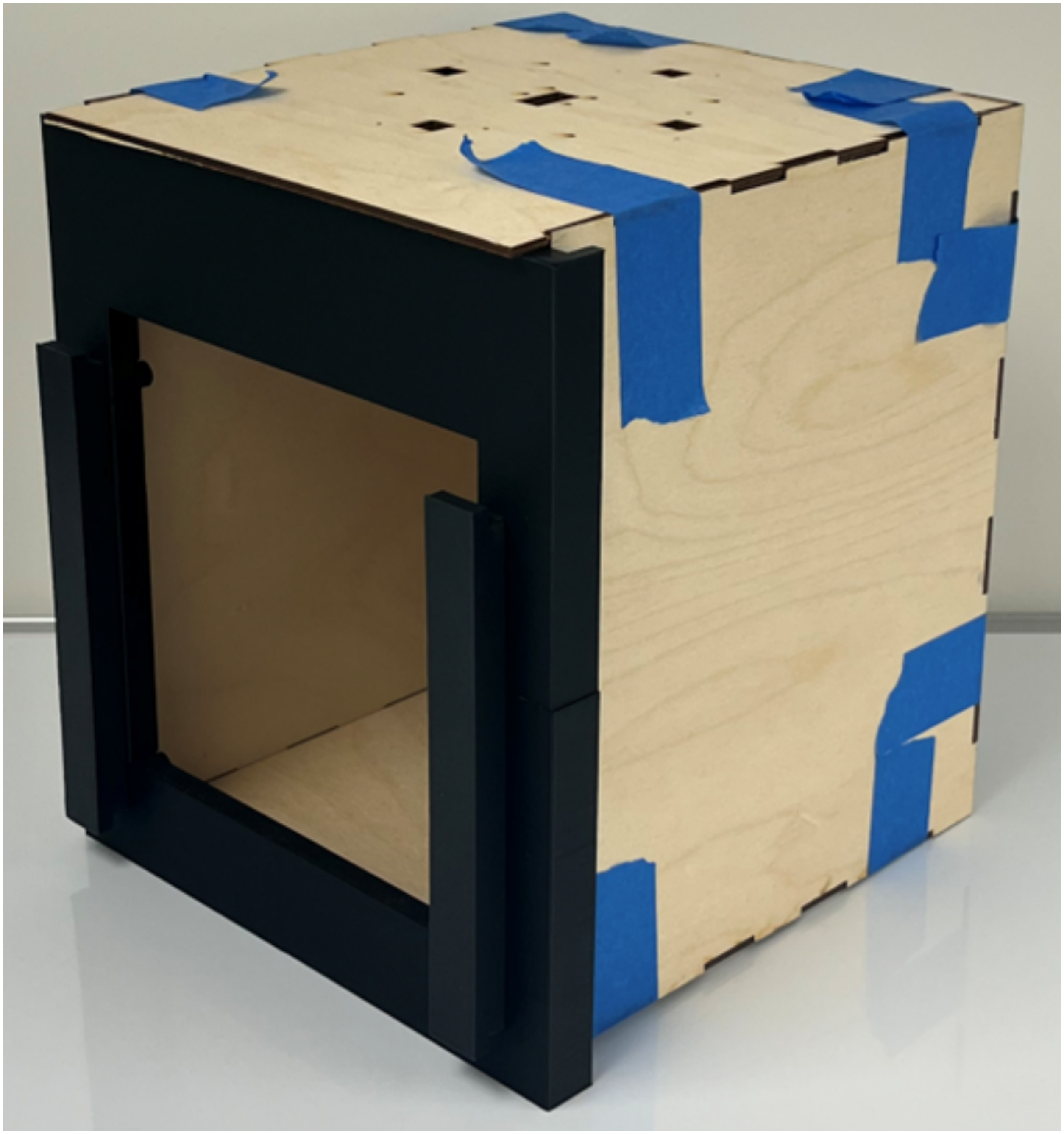

**SFig 13 Imaging box from interlocking wood panels**

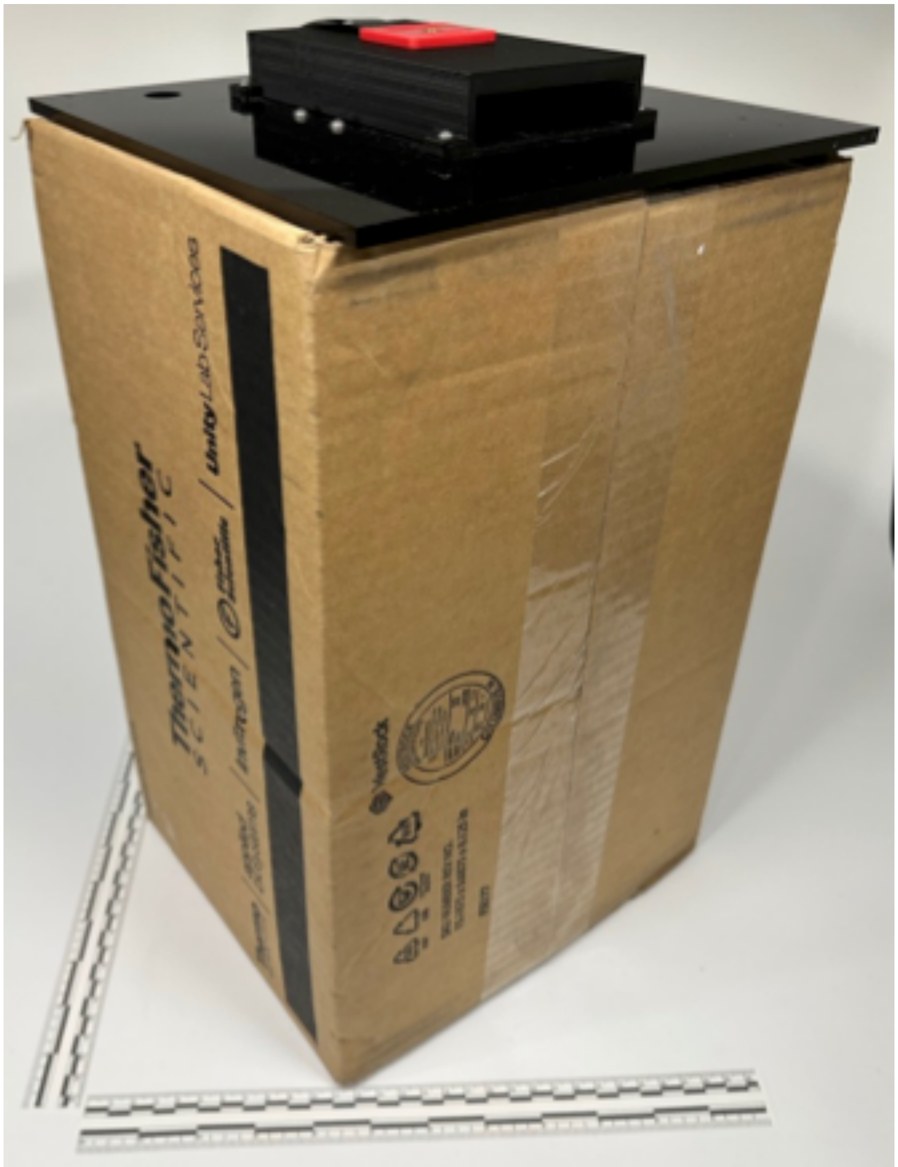


**SFig 14 Imaging box from standard cardboard box**

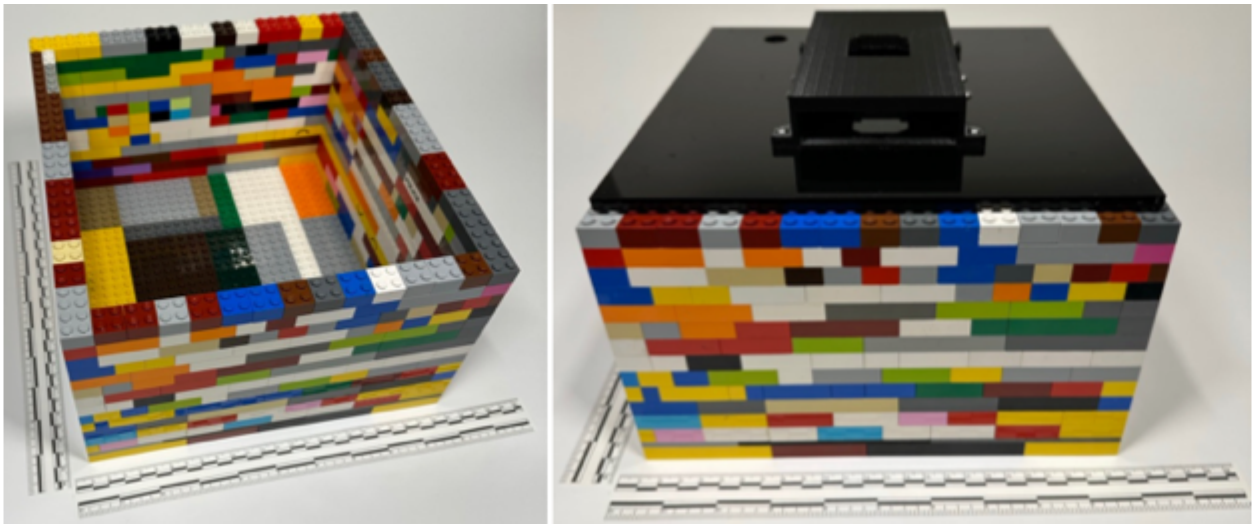

**SFig 15 Imaging box from standard Lego pieces**

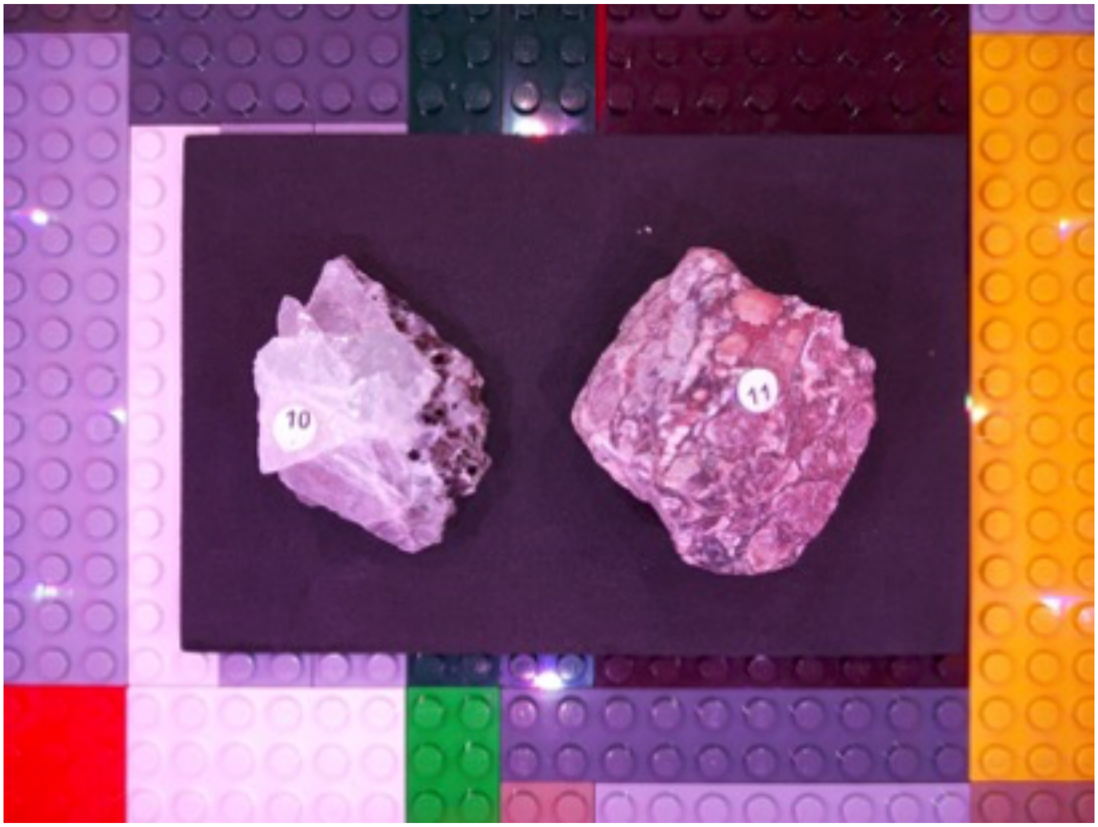


**SFig 16 Image of minerals in Lego imaging box using white LEDs** Left mineral is Fluorite from Utah, Right mineral is Turritella Agate from Wyoming
